# Waveform morphology links local circuit kinetics to cortical interactions

**DOI:** 10.64898/2026.11.13.766469

**Authors:** Pablo Núñez, Gulshan Kumar, Clarisse Gotti, Peng Wang, Laurence Martineau, Paule Lessard Bonaventure, Roxane Hoyer, Bratislav Mišić, Philippe Albouy, Steven Laureys, Prejaas K.B. Tewarie

**Author notes:** These authors contributed equally to this work (Pablo Núñez, Gulshan Kumar).

## Abstract

How does local cortical physiology shape interactions between brain regions? We combined neural mass modelling with human and macaque intracranial recordings to examine how synaptic kinetics link oscillatory waveform morphology to cortical interactions. In a modified excitatory–inhibitory Wilson–Cowan model, synaptic rise and decay kinetics shaped waveform geometry and phase-dependent responsiveness. Local kinetics and coupling symmetry jointly influenced interaction measures and temporal irreversibility, while phase-locked perturbations altered imaginary coherency and bivariate irreversibility even when the coupling strengths remained fixed. In stereo-EEG from 23 patients, waveform asymmetry was most strongly associated with local irreversibility, whereas mean instantaneous alpha frequency showed stronger associations with interregional measures. Subdural macaque electrocorticography reproduced the frequency associations but not the local wave-form association. Together, these findings show that waveform shape carries information about local neuronal dynamics, provide a mechanistic framework for interpreting cortical interactions, and motivate testing waveform morphology as a guide to stimulation timing.

---

Neuronal oscillations organise the timing of communication between cortical regions: the effect of an incoming signal depends partly on the moment when it reaches the receiving population [1]. Excitatory and inhibitory populations have synaptic responses that rise and decay over different time scales, contributing to the temporal structure of their activity, and these can differ across different states and populations. Receptor dynamics, excitation–inhibition interactions and recurrent activity all contribute to this temporal variability [2, 3, 4]. Its effects may be visible by the naked eye in the recorded signal, whose rhythms can rise sharply and decay slowly, have broad peaks and narrow troughs, or approach a sinusoid, even at the same frequency [5, 6]. In this study, we asked whether changing the local dynamics expressed in a neuronal waveform can also change the interactions measured between regions when their internal coupling is held fixed.

Interpreting neural activity through its temporal shape goes back a long way. In particular, studies on event-related potential distinguish components by the amplitudes and latencies of peaks in responses aligned to a stimulus [7, 8, 9]. Waveform morphology and its variability has been linked to local physiology and indeed several studies have found that it carries physiological information. For instance, beta sharpness in human motor cortex is associated with Parkinsonian pathophysiology [10], the shape of transient neocortical beta events reflects the relative timing of proximal and distal synaptic drives in detailed models and laminar recordings [11], and within-cycle instantaneous frequency resolves the temporal structure of individual oscillations [12]. Within-cycle instantaneous frequency provides another description of this temporal geometry [12]. These observations motivate waveform morphology as an important element of local dynamics, but what it specifically tells us about interactions between cortical regions still needs to established.

From another perspective, interregional phase relationships, coherence and directed spectral measures describe different aspects of coordinated brain activity [13, 14, 15]. Crucially, these measures are obtained from narrowband-filtered signals, and this filtering forces each cycle towards a sinusoidal shape, so the within-cycle structure that the aforementioned literature showed as informative is eliminated before the interaction is estimated in the first place. [16, 17]. Temporal irreversibility provides another point of view, by asking whether the dynamics of the activity differ when examined forward and backward in time [18, 19, 20, 21]. Because both the local generator within a population and the coupling between brain regions shape the observed signals, a change in an interaction measure does not necessarily identify which of them changed.

We hypothesised that these two interpretations of neural activity are likely not independent, because the same synaptic kinetics could underlie both. A change in a neuronal population’s kinetics alters the waveform it generates and transmits but, at the same time, also alters the window over which the population integrates incoming activity, with dendritic processing and recurrent connections further shaping this response [22, 23]. In this study, we tried to reconcile these two perspectives by means of a a modified Wilson–Cowan excitatory–inhibitory neural mass model whose effective synaptic rise and decay time scales can be varied while interregional coupling is controlled [24, 25]. We examined fluctuations near a steady state and in a sustained oscillation regime, measuring how local kinetics can affect waveform geometry, population interactions, temporal irreversibility and responses to stimulation at different phases. We then analysed real human stereo-EEG and macaque electrocorticography to establish whether measurable waveform features covary with local and interregional statistics in cortical recordings, using masked empirical mode decomposition to quantify within-cycle morphology [26, 12].

We found that that local kinetics jointly shape waveform geometry and susceptibility in the model, while coupling between regions and population input determine how the directional imbalance is expressed. Using mixed-effects models, human recordings showed correlations between waveform and local temporal irreversibility, and a stronger associations between instantaneous frequency and interregional measures. The frequency associations replicated in macaque recordings, but the local waveform association did not. These findings provide a mechanistic account of how local temporal dynamics enter cortical interactions and identify the conditions under which waveform morphology may help interpret those interactions. We conclude that the information about local temporal dynamics carried by waveform morphology should be considered in conjunction with measures of interregional interaction and irreversibility.

## Results

We used a modified Wilson–Cowan neural mass model to study whether local synaptic kinetics could affect waveform morphology, local irreversibility and its response to timed perturbations. In addition, we studied how measures of interaction, network irreversibility, and perturbation propagation are affected when the underlying coupling between regions remains fixed. In the model simulations, each node represents a local cortical region, with an excitatory and an inhibitory population. Their firing rates drive synaptic responses whose effective rise and decay time scales can be set separately (as opposed to traditional Wilson–Cowan models), and the filtered excitatory output is used as the proxy for the observed signal (i.e., the signal recorded in an EEG). The excitatory output also provides input to the excitatory population of another node (or multiple nodes, in network simulations). The coupling strength can also be set independently in each direction between pairs of nodes. We examined two distinct operating regimes: the first one was near the onset of oscillation, where noise produces fluctuations around a steady state, allowing us to study transmission and spectral connectivity. The second regime was that of sustained cycles, which allowed us to study waveform geometry, irreversibility and phase-dependent responses. We first examined one node, then pairs and networks of nodes, and finally tested relationships among observable features in intracranial recordings from humans and macaques.

### Synaptic kinetics shape local waveform geometry

We first investigated whether changing synaptic kinetics in the model could alter the waveform generated by one local node of the modified model. As expected, when external excitatory drive increased, the node moved from a stable state to a sustained oscillation at approximately 4– 5 Hz to a supercritical Hopf bifurcation. This showed that we could use a controlled rhythm in which to vary synaptic decay while holding the other model parameters fixed for the remaining tests. With rise times fixed, shortening synaptic decay produced sharper, more asymmetric cycles, while longer decay attenuated higher harmonics and made the waveform more sinusoidal (Figure 1b–d). Measured rise and decay durations became more similar as the synapse slowed, and rise–decay symmetry approached 0.5. We thus showed that synaptic decay was able to control waveform asymmetry in our model.

**Figure 1:**
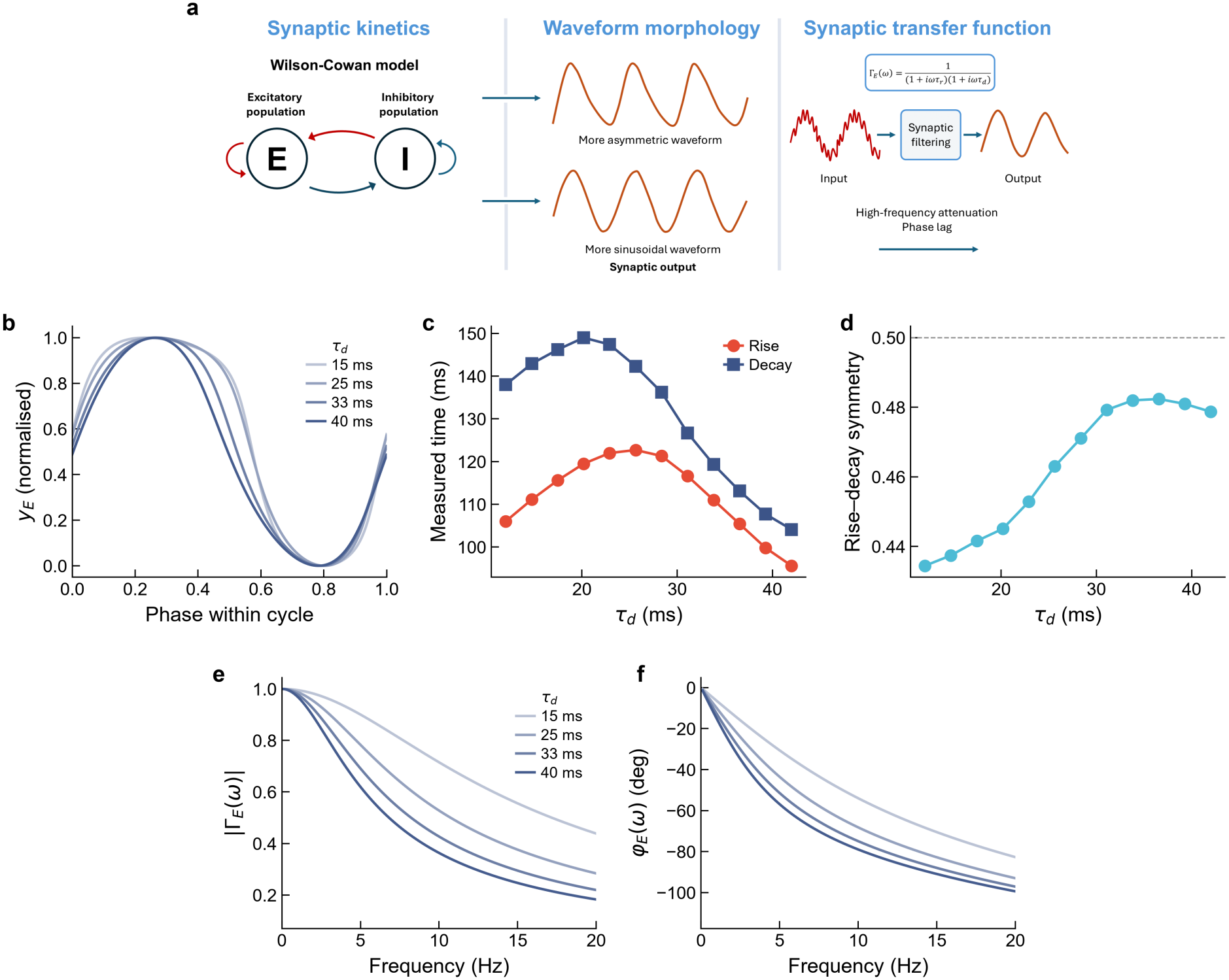
The dual-exponential synapse controls waveform morphology through its complex transfer function. (a) Schematic overview of an excitatory–inhibitory circuit with dual-exponential synaptic integration. With rise time fixed, shorter and longer decay times yield different synaptic responses and oscillatory waveform shapes. (b) One normalised cycle of the excitatory synaptic output for increasing synaptic decay times *τ*_*d*_ with fixed rise times. A short decay displays a sharper, more asymmetric waveform, whereas increasing *τ*_*d*_ pushes the activity closer to that of a sinusoid. (c) Cycle-by-cycle rise and decay times measured from the simulated waveform as a function of *τ*_*d*_. The two are most unequal for short decay times and converge as *τ*_*d*_ increases. (d) Rise–decay symmetry (where 0.5 indicates a symmetric waveform) as a function of *τ*_*d*_. All waveforms were obtained in a supercritical self-oscillating regime. (e) Magnitude |Γ_*E*_(*ω*)| of the synaptic transfer function for different synaptic decay times *τ*_*d*_: larger *τ*_*d*_ produces stronger low-pass attenuation. (f) Phase *φ*_*E*_(*ω*) of the transfer function for the same decay times.

The synaptic transfer function within the model Γ_*E*_(*ω*) describes how synaptic filtering changes the amplitude and phase of an input at each frequency in the model. Its magnitude determines attenuation at each frequency, while its phase describes the associated lag. Increasing decay time attenuated high frequencies more strongly and increased the lag (Figure 1e,f). Both effects are present even when rise and decay time constants are equal. Within the recurrent circuit model, these kinetics influence the outgoing waveform and the response to incoming activity. The overall response also depends on the strength of feedback within the local circuit and its baseline level of activity.

### Reciprocal interaction between populations does not follow underlying directional imbalances

Next, we investigated how local kinetics have an effect on the interaction between two model populations. To this end, we compared linear-response predictions with stochastic simulations of two weakly coupled nodes near the fixed point, averaging spectra across 30 runs. Coherence was nonzero for both reciprocal and unidirectional coupling and varied with synaptic filtering (Figure 2b). Imaginary coherency cancelled out in the identical, reciprocally coupled pair driven by equal independent noise. Unidirectional coupling broke the exchange symmetry and produced a nonzero imaginary component whose frequency dependence reflected local kinetics (Figure 2c). Across additional supercritical operating points, symmetric-pair imaginary coherency remained small relative to the directed condition (Supplementary Figure S2).

**Figure 2:**
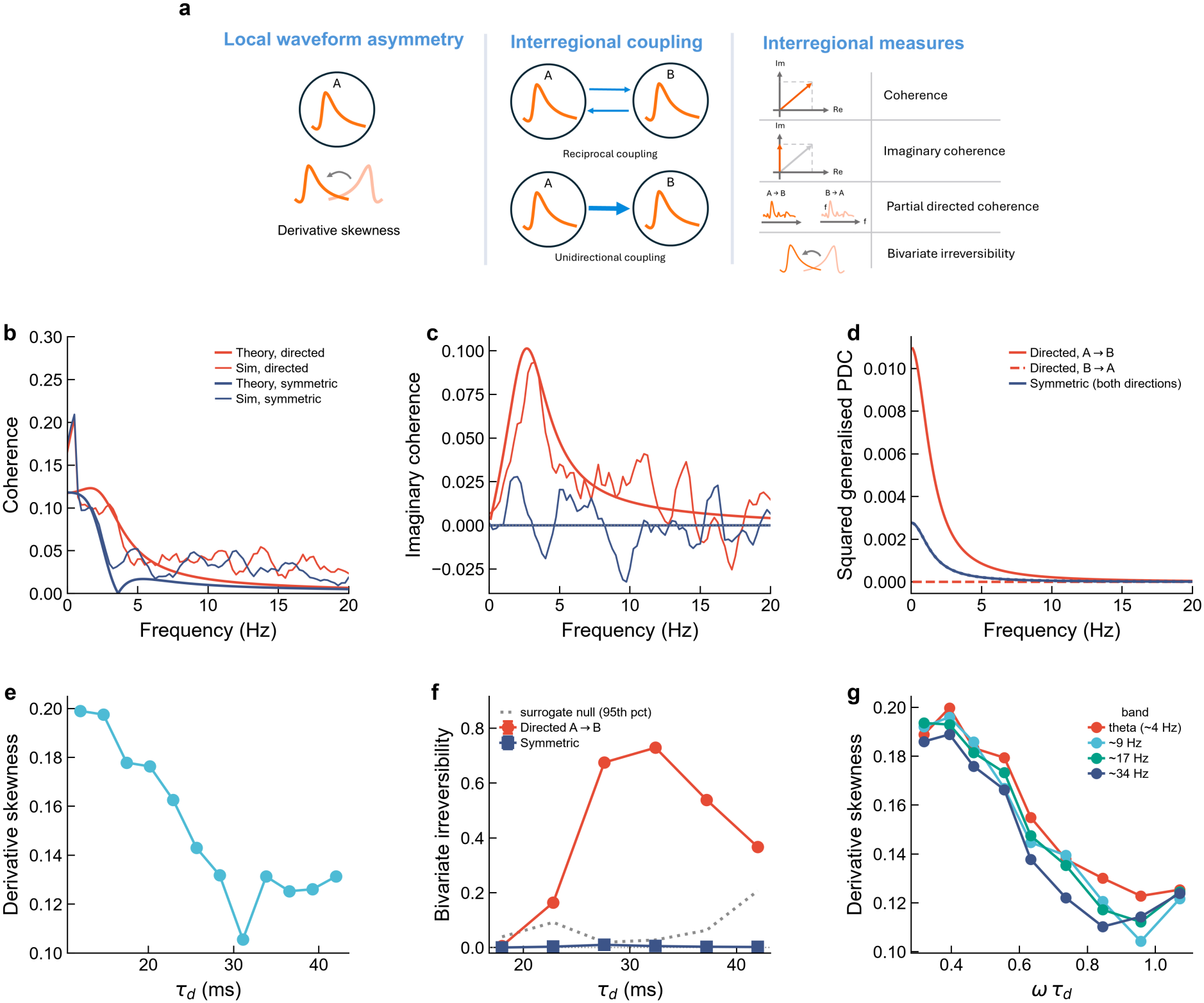
Local kinetics affect connectivity (b–d) and irreversibility (e–f). (a) Schematic distinction between local waveform asymmetry and interregional measures in two nodes with identical local kinetics. Symmetric reciprocal and unidirectional coupling are compared at matched total coupling strength. Derivative skewness describes local temporal asymmetry, whereas coherence, imaginary coherency, squared generalised partial directed coherence (gPDC) and bivariate irreversibility describe distinct aspects of between-node dynamics. (b-d) Two–node connectivity. Analytical linear-response predictions (thick lines) are compared against ensemble stochastic simulations (thin lines), for directional A→B coupling (red) and symmetric coupling (blue), at matched total coupling strength. (b) Coherence magnitude is observable for both coupling configurations and is shaped by |Γ_*E*_(*ω*)|. (c) For the identical, equally and independently driven pair, imaginary coherency is theoretically zero under reciprocal coupling and near zero in simulation (blue), with a nonzero value under unidirectional coupling (red). (d) Squared generalised partial directed coherence (computed analytically) for the directed configuration (red: A→B solid, B→A dashed) and for the symmetric configuration (blue). (e) Univariate time-irreversibility (derivative skewness) as a function of the synaptic decay time. (f) Bivariate time-irreversibility (forward-versus-reversed shifted-correlation measure) for symmetric versus directional coupling, as a function of *τ*_*d*_, measured in the supercritical regime, across 30 independent noise realisations per condition and synaptic decay time. The dotted line is the 95th percentile of independent Fourier phase-randomised surrogates, which preserve individual spectra but eliminate interchannel phase relationships. (g) Derivative skewness across four oscillation bands (theta to gamma, obtained by uniform time-rescaling of the model), plotted against the dimensionless product *ωτ*_*d*_.

Analytical squared generalised partial directed coherence (gPDC, a measure of connectivity that characterises the directed linear relationship between simultaneously observed time series in the frequency domain) [27] was nonzero and equal in both directions under underlying reciprocal coupling between populations. Unidirectional coupling produced a directional imbalance, modulated by synaptic filtering (Figure 2d). Reciprocal coupling therefore supported influence in both directions while preserving equality between their analytical PDC values.

We observed that autoregressive fits followed the broad frequency dependence of analytical A→B gPDC, but their magnitude varied with model order. They also produced a nonzero B→A estimate when the reverse connection was absent, approaching the forward estimate at higher frequencies (Supplementary Figure S1). We therefore used analytical gPDC to compare the coupling configurations.

### Local and interregional temporal asymmetry have different dynamics

We then studied whether an asymmetric local waveform leads to an asymmetric relationship between populations. We did so by comparing local and bivariate irreversibility, which quantify how much the activity of individual populations and pairs of populations differs when viewed forward and backward in time. At a single node, derivative skewness changed with synaptic decay time and approached zero as the waveform became more sinusoidal (Figure 2e). Uniformly rescaling the model time constants from theta to gamma preserved the relationship when expressed against *ωτ*_*d*_ (Figure 2g). This collapse under time rescaling confirmed the role of the synaptic time scale relative to the oscillation period within our model.

The bivariate index compares transformed lagged correlations before and after time reversal [20, 21]. In the model pairs it was negligible for reciprocal coupling but appreciable for unidirectional coupling, with a non-monotonic dependence on synaptic decay (Figure 2f). We thus showed that local waveform asymmetry can coexist with negligible coupling irreversibility when the underlying coupling is symmetric.

### Network organisation modulates global irreversibility

We extended the analysis to networks of neural mass models with varying reciprocity, synaptic rise–decay ratio and size (Figure 3). Across 60 independent realisations, we observed that lower reciprocity between nodes increased global irreversibility most clearly in the smallest networks. Symmetric networks also showed a nonzero baseline, demonstrating that the null observed in an exchange-symmetric pair does not necessarily carry over to a larger nonlinear network. This showed that even equally coupled populations can behave differently because they receive different inputs from the surrounding network.

**Figure 3:**
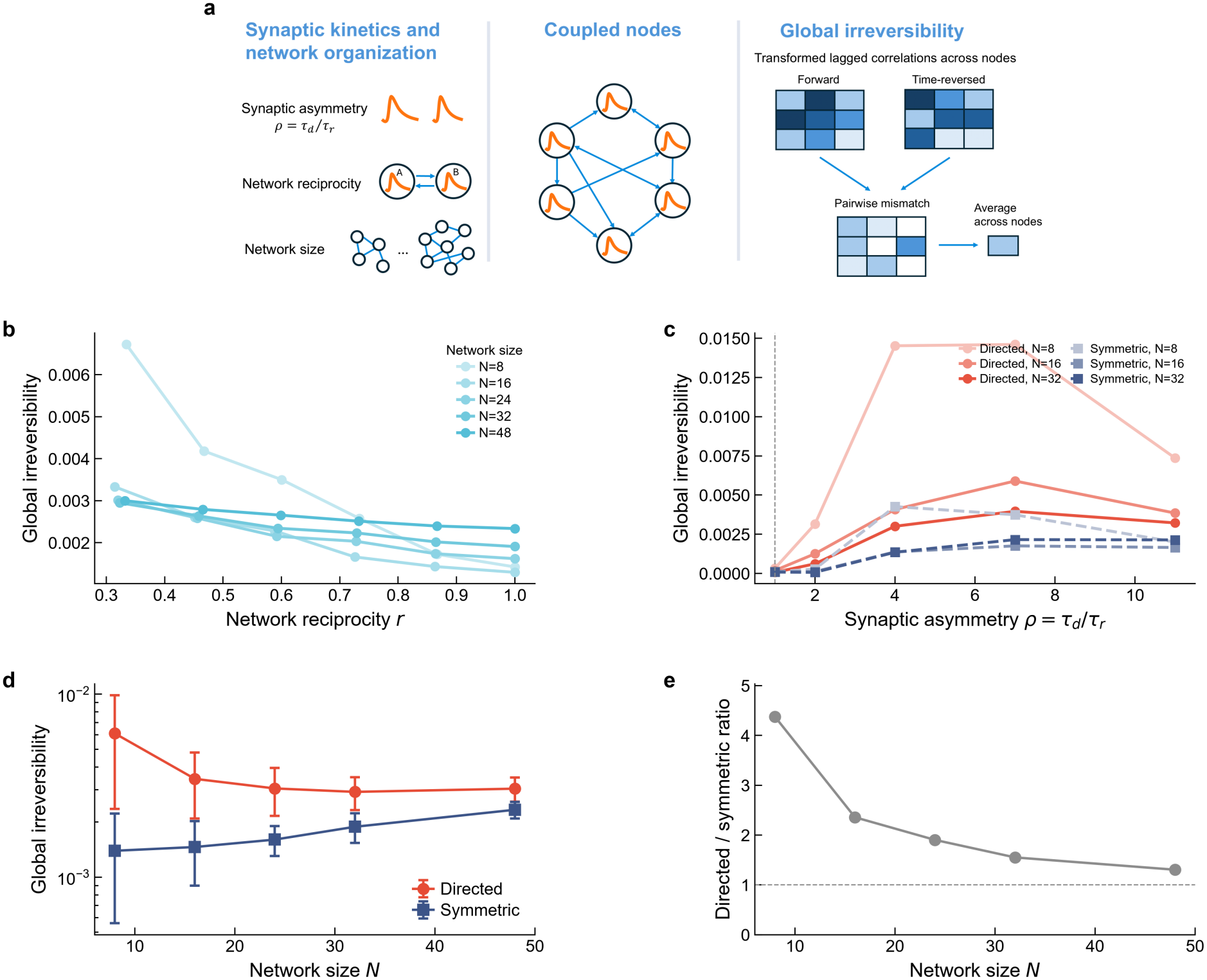
Synaptic kinetics and network organisation modulate irreversibility together. (a) Schematic overview of synaptic asymmetry, network reciprocity (*r*) and network size (*N*) in coupled oscillating nodes. Global irreversibility is obtained by comparing transformed lagged correlations from forward and time-reversed activity and averaging their squared differences across node pairs. (b) Global time-irreversibility as a function of network reciprocity *r* at matched per-node input strength, repeated across network sizes *N* ∈ {8, 16, 24, 32, 48} (light to dark). (c) Global irreversibility as a function of synaptic asymmetry *ρ* = *τ*_*d*_*/τ*_*r*_, for symmetric (dashed) and directed (solid) networks at network sizes *N* ∈ {8, 16, 32} (light to dark). (d) Global irreversibility as a function of network size *N*, for symmetric and directed networks. (e) Ratio of the directed and symmetric mean global irreversibility as network size increases. In (b)–(d), markers denote means across 60 independent network realisations per condition. Panel (d) shows standard-deviation error bars. Error bars are omitted in (b) and (c) for clarity.

Varying the synaptic asymmetry ratio *ρ* = *τ*_*d*_*/τ*_*r*_ at fixed geometric-mean time scale showed an inverted-U dependence: the index was smallest near *ρ* = 1, peaked near *ρ* ≈ 4, and declined at larger ratios (Figure 3c), with the effects again being more marked the smaller the network. Repeating the global irreversibility sweep at networks of size *N* = 8, 16, 24, 32, 48 showed a progressively flatter dependence on size, whereas the synaptic-ratio sweep retained its qualitative shape at *N* = 8, 16, 32 (Figure 3d,e). These results show that local kinetics and network organisation jointly affect global irreversibility in the networks of neural mass models.

### Local kinetics shape phase-dependent responses and their propagation over node couples

We perturbed the excitatory rate of a node at different phases of its ongoing cycle to test for the consequences of timed perturbations on connectivity and irreversibility in single nodes and node pairs. For a fast synapse, the phase-response curve had a sharp positive peak near the rising flank and a broader negative valley later. For a slower synapse, the response was smoother and its positive peak occurred earlier (Figure 4b). The model kinetics therefore changed both the observed waveform and the phases at which a pulse most effectively shifted the model’s timing.

**Figure 4:**
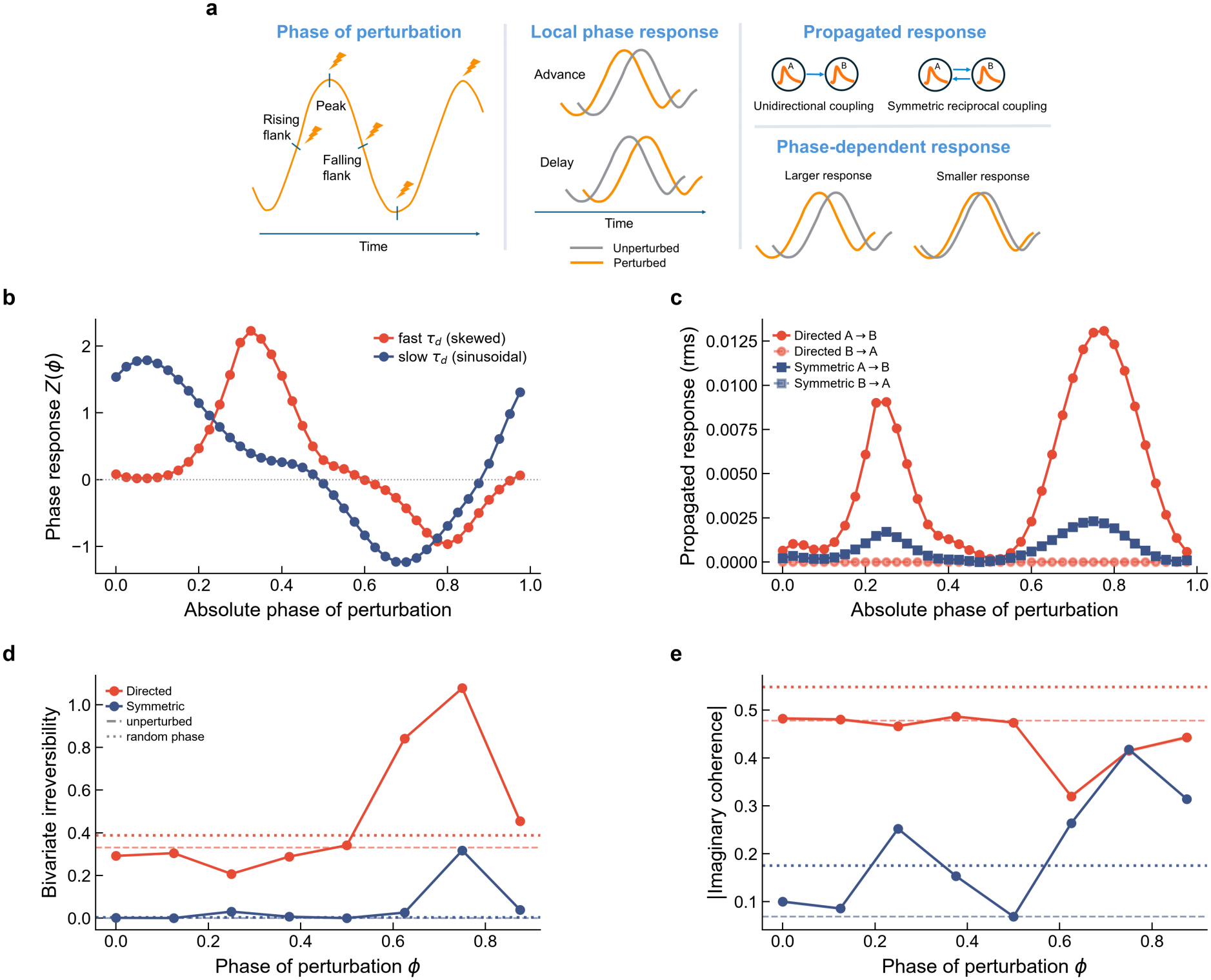
The timing of a perturbation shapes the connectivity and irreversibility it induces. (a) Schematic overview of phase-dependent perturbation and propagation. A pulse delivered at different oscillation phases can advance or delay local activity and change the downstream response. (b) Single-node phase response: the phase shift produced by a small pulse to the excitatory rate variable, as a function of the absolute phase at which it is delivered, for a fast, skewed synapse (*τ*_*d*_ = 15 ms) and a slow, near-sinusoidal synapse (*τ*_*d*_ = 40 ms). (c) Two coupled nodes: root-mean-square propagated response of a pulse delivered to A and read out at B (A→B) and the reverse (B→A), against perturbation phase, for unidirectional (directed) and for matched-total-strength reciprocal (symmetric) coupling. Propagation is one-directional for the directed configuration For the symmetric configuration, the A→B and B→A curves coincide. (d) Noisy coupled nodes: bivariate time-irreversibility as a function of the phase of a perturbation train applied to a single node, for directed and symmetric coupling, averaged over *n* = 30 independent noise realisations. The dashed lines mark the unperturbed baselines. (e) The same for the band-averaged magnitude of imaginary coherency. Panels (c)–(e) use a fast synapse, (c) is deterministic and (d, e) are noise-driven near the oscillatory onset.

With two coupled nodes, an upstream pulse propagated downstream under unidirectional coupling, while a downstream pulse had no upstream effect, and the strongest perturbations where found to follow the cycles peaks and valleys. Symmetric coupling produced equal propagated responses in the two directions, each strongly dependent on stimulation phase (Figure 4c). The larger response along the unidirectional pathway can be interpreted in light of that of the matched total coupling: the unidirectional configuration concentrated the weight on one edge, and the reciprocal configuration divided it between two edges.

In noisy model node pairs, phase-locked perturbative pulse trains applied to a single node changed bivariate irreversibility and imaginary coherency differently across the oscillatory cycle (Figure 4d,e). Precisely timed stimulation on the models could produce nonzero values in a structurally symmetric pair, and imaginary coherency in the directed pair could decrease at some phases. These results demonstrate a way in which the timing of a local input can alter interactions between nodes without a change in the underlying structural coupling being present.

### Waveform morphology and instantaneous frequency predict irreversibility and connectivity in human recordings

The modified Wilson–Cowan model showed how local kinetics can affect several observables under controlled coupling. However, it is not feasible to isolate said kinetics in electroencephalo-graphic recordings, since the “real” underlying coupling between regions cannot be accurately estimated. Therefore, for the human and macaque recordings we tested the following prediction: whether differences in the shape and frequency of an ongoing rhythm are associated with local and between-channel metrics. We focused on the alpha frequency band, as it could be successfully identified in both intracranial datasets and its cycles could be measured individually.

In total, we analysed 5,602 stimulation runs from 23 patients (25 recording sessions). The bipolar montage included 2,205 grey-matter channels (median 97 per patient, range 58–141), yielding 320,255 post-stimulation channel-level observations. The alpha mode had a median instantaneous frequency of 9.98 Hz (interquartile range 9.48–10.50 Hz). Median ascent–descent and peak–trough ratios were 0.506 and 0.490, respectively (Figure 5).

**Figure 5:**
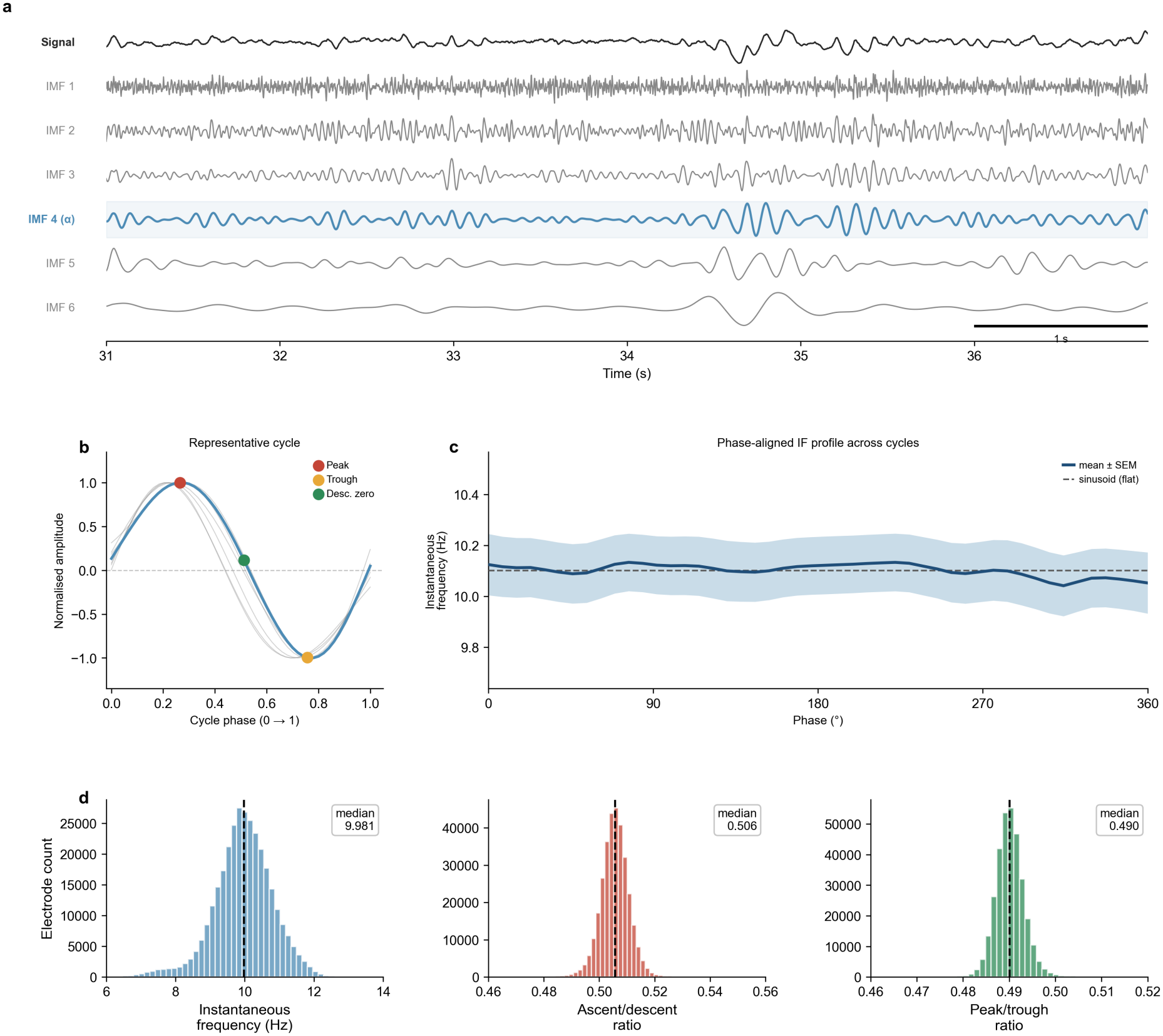
Isolating the alpha rhythm and quantifying cycle shape. (a) A 6-s excerpt of a post-stimulation bipolar recording (top) and the six intrinsic mode functions returned by masked empirical mode decomposition. The fourth mode, shaded, corresponds to the 10-Hz mask and carries the alpha rhythm. (b) One cycle of that mode, normalised in amplitude and duration, with the peak, trough and descending zero crossing marked. Grey traces show other cycles from the same electrode. The relative timing of these landmarks defines the ascent-to-descent and peak-to-trough ratios. (c) Mean phase-aligned instantaneous-frequency profile across all retained cycles at the same electrode, shaded ± SEM, against a flat profile (dashed). This electrode was selected as closest to the population medians of the three waveform features and is therefore near-symmetric. (d) Distributions of the three waveform features across the 320,255 channel-level observations in the dataset, with medians marked.

In mixed-effects models, 14 of the 15 coefficients across mean frequency and the two waveform morphology ratios survived multiple-comparison correction. Ascent–descent ratio was negatively associated with coherence, imaginary coherency, PDC asymmetry and univariate irreversibility (all FDR-corrected *p <* 0.001), but not bivariate irreversibility (*β* = +0.001, *p* = 0.66, Figure 6). Its strongest association was with univariate irreversibility (*β* = −0.107).Mean frequency showed a complementary pattern: larger effects on the four between-channel out-comes (*β* = −0.035 to −0.126), with a smaller positive coefficient for univariate irreversibility (*β* = +0.017). The unadjusted rank correlation between ascent–descent ratio and univariate irreversibility was −0.347. Its magnitude did not exceed 0.050 for other outcomes (Supplementary Figure S4).

**Figure 6:**
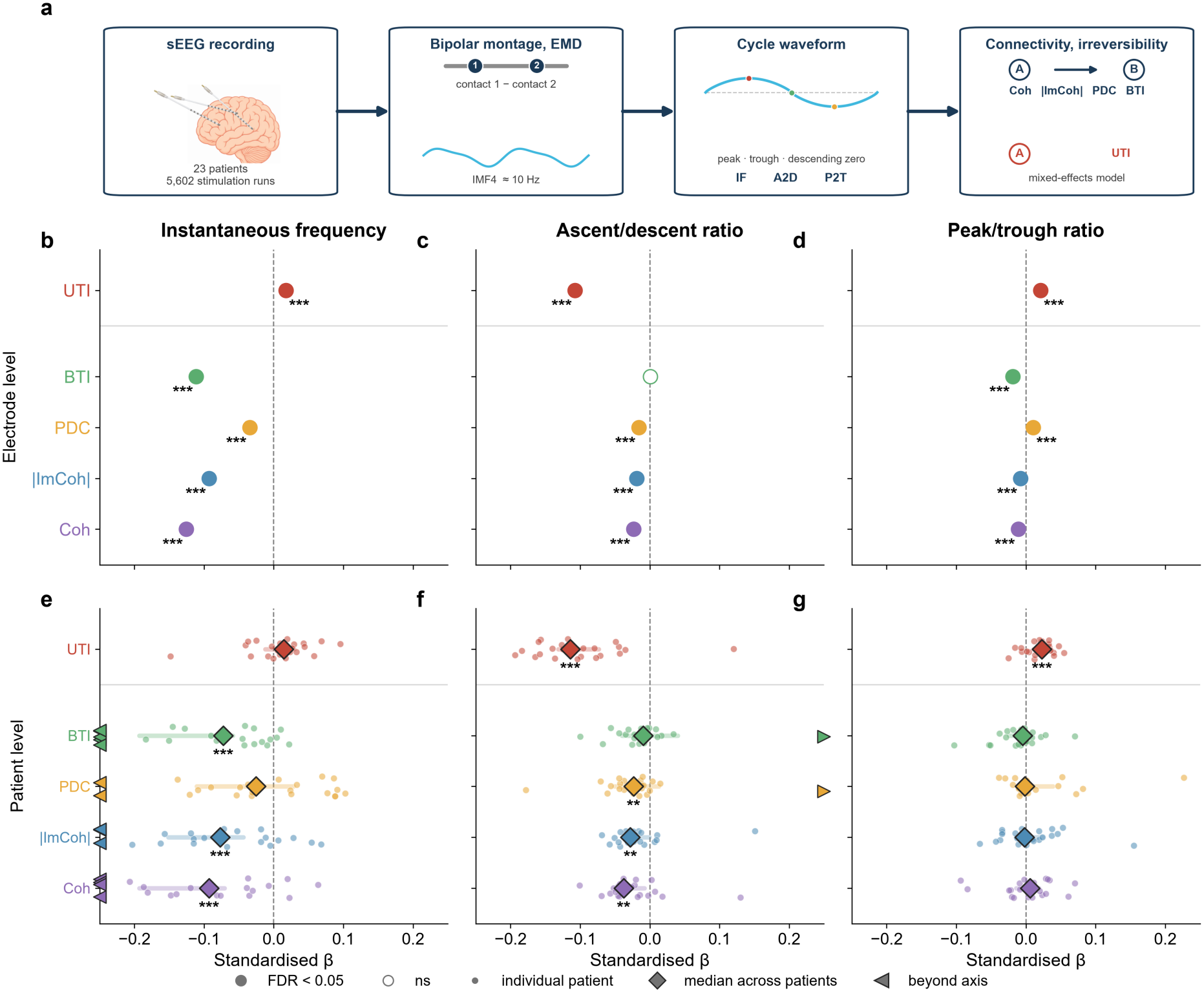
Waveform shape and instantaneous frequency predict different measures in human sEEG. (a) Analysis pipeline. Stereo-EEG recordings from 23 patients during focal stimulation were re-referenced to a bipolar montage and decomposed using masked empirical mode decomposition. The fourth mode carried the alpha rhythm. Each cycle was summarised by its mean instantaneous frequency (IF) and two control-point ratios: ascent-to-descent (A2D) and peak-to-trough (P2T). Standardised coefficients were obtained from linear mixed-effects models, one per outcome, controlling for brain region, stimulation protocol and stimulated-channel status. Columns show the three waveform features, and the five outcomes are stacked within each panel, with univariate time-irreversibility separated from the four between-channel measures by a horizontal line. (b–d) Channel-level analysis, including all observations with patient as a random effect. Filled markers indicate FDR-corrected *p <* 0.05, whereas open markers indicate non-significant effects. Stars show the FDR-corrected significance of the mixed-model coefficients: *** *p <* 0.001, ** *p <* 0.01, * *p <* 0.05; ns, not significant. The 95% confidence intervals are shown but are narrower than the markers at this sample size. (e–g) The same model fitted separately within each of the 23 patients. Small dots represent individual patients, diamonds show the median across patients, and horizontal bars show the 95% confidence interval. Triangles identify patients whose slopes extend beyond the displayed axis. Stars show FDR-corrected one-sample Wilcoxon signed-rank tests. Coh, coherence. |ImCoh|, magnitude of imaginary coherency. PDC, partial directed coherence asymmetry. BTI, bivariate time-irreversibility. UTI, univariate time-irreversibility.

Patient-level inference supported the main dissociation: 8 out of 15 effects survived correction when within-patient coefficients were tested across 23 patients. The ascent–descent association with univariate irreversibility had the same sign in 22 patients (median *β* = −0.114, FDR-corrected *p <* 0.001), whereas its frequency association was not significant. Instantaneous frequency associations with coherence, imaginary coherency and bivariate irreversibility remained significant.

Among between-channel outcomes, coherence had the largest adjusted association with ascent– descent asymmetry at both channel and patient levels (FDR-corrected *p <* 0.01). This ordering was absent from unadjusted correlations (Supplementary Figure S6). Partial regressions for the primary models are shown in Supplementary Figure S7. We also measured how strongly instantaneous frequency varied within a typical alpha cycle, using the range of the phase-aligned mean frequency profile divided by its mean. This measure remained associated with coherence (*β* = +0.050), imaginary coherency (*β* = +0.026), PDC asymmetry (*β* = −0.029) and bivariate irreversibility (*β* = +0.085) after adjustment for mean frequency and the two morphology ratios (all FDR-corrected *p <* 0.001).

At the electrode-pair level, larger differences in mean instantaneous frequency were associated with slightly lower coherence (*ρ* = −0.054, permutation *p <* 0.001). Associations with imaginary coherency were smaller under both stimulation protocols (*ρ* = −0.024 at 1 Hz and −0.028 at 50 Hz). The signed log ratio of the electrodes’ ascent–descent ratios was also weakly associated with their directional PDC imbalance (*ρ* = −0.016 in 23 patients at 1 Hz and −0.013 in 23 patients at 50 Hz, both *p <* 0.001). These correlations exceeded those obtained after waveform values were shuffled among electrodes while the connectivity matrices were held fixed, but their magnitudes were small.

Stimulated contacts showed larger changes in both shape ratios than the other contacts (both *p <* 0.001, Supplementary Figure S5). However, changes in either ratio were not associated with changes in connectivity. Frequency changes were associated with connectivity changes, most clearly PDC (*ρ* = 0.179, *p <* 0.001).

### Macaque recordings reproduce the instantaneous frequency associations found in humans but not the local waveform associations

We repeated the analysis in awake, eyes-closed electrocorticography from four macaques, comprising 31 sessions and 1,431 bipolar channel–session observations. In these recordings the alpha mode was centred at 11.5 Hz. Preprocessing was adapted to the recording geometry and mains frequency, as well as statistical grouping used session.

Mean instantaneous frequency was negatively associated with coherence (*β* = −0.344), imaginary coherency (*β* = −0.229), bivariate irreversibility (*β* = −0.224) and PDC asymmetry (*β* = −0.069, Figure 7). Coherence had the largest and PDC the smallest frequency effect in both datasets. Within-session frequency effects on coherence, imaginary coherency and bi-variate irreversibility survived correction, with negative slopes in 26, 27 and 30 of 31 sessions, respectively.

**Figure 7:**
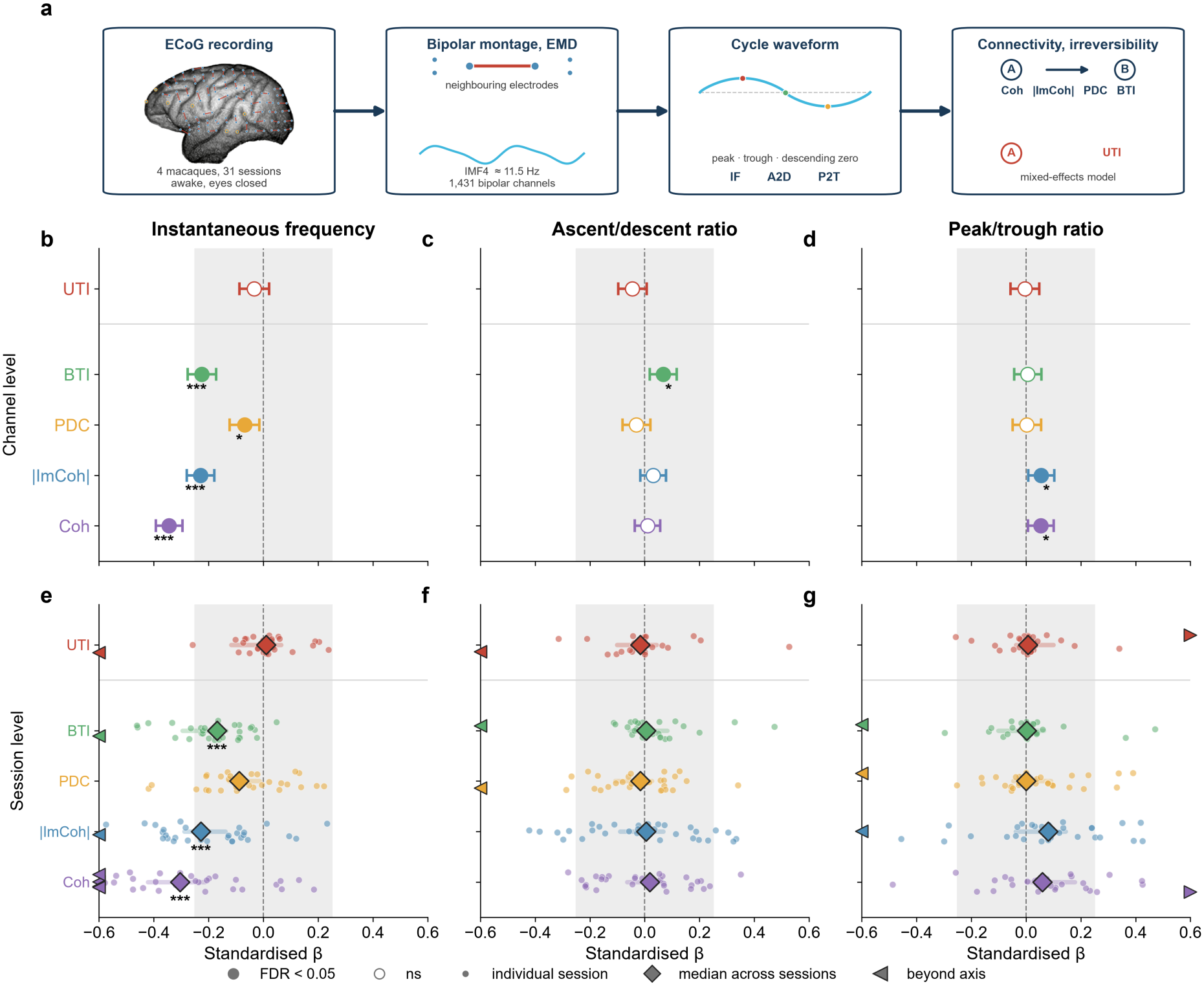
Waveform shape and instantaneous-frequency associations in macaque electrocorticography. (a) Analysis pipeline.Electrodes were paired into bipolar channels using minimum-distance matching across the array surface, and the fourth intrinsic mode carried an alpha rhythm centred at 11.5 Hz. The analysis included 1,431 bipolar channels from 31 awake, eyes-closed sessions in four macaques, with session replacing patient as the random effect and monkey included as a fixed covariate. (b–d) Channel-level results. (e–g) Session-level results. The shaded band marks the axis range used in Figure 6, allowing comparison between the human and macaque results. Coh, coherence. |ImCoh|, magnitude of imaginary coherency. PDC, partial directed coherence asymmetry. BTI, bivariate time-irreversibility. UTI, univariate time-irreversibility.

The association between ascent–descent ratio and univariate irreversibility found in humans did not replicate (*β* = −0.045, *p >* 0.05). Although its sign was negative in 21 sessions, no waveform effect on this outcome survived correction at channel or session level.

## Discussion

In the present study we investigated how the physiology expressed in a local cortical oscillation shapes interactions between populations. In the dual-exponential modified Wilson–Cowan neural mass model, changing local synaptic kinetics altered measures of interaction between populations, with the coupling weights between nodes were fixed, thus showing that a change in a network statistic can arise simply from a change in the waveform kinetics of the populations involved. The human and macaque recordings allowed an initial test of correlations among measurable features. In the human cohort, the clearest shape association was with a local measure of temporal asymmetry, whereas between-region measures showed larger associations with alpha instantaneous frequency. However, the local waveform shape association did not replicate in the macaque population.

In the model, each node sends a synaptically filtered version of its excitatory activity to another node. Synaptic rise and decay times change the shape and timing of this outgoing signal. Its effect depends on when it reaches the receiving node, whose sensitivity to input varies across the oscillation cycle [1]. The phase-interaction function combines the incoming waveform with the receiving node’s phase response and local gain to describe its influence at each relative phase. Synaptic kinetics can therefore change when and how strongly connected nodes influence one another even when their coupling strength is fixed.

Our model captures finite synaptic integration within a recurrent excitatory–inhibitory circuit. In biological tissue, receptor dynamics, dendritic filtering and the timing of incoming activity can also shape population responses [2, 22, 23]. For example, the relative timing and strength of proximal and distal inputs contribute to the shape of transient neocortical beta events [11]. Excitation and inhibition are also coordinated within individual hippocampal gamma cycles [4]. Whether these different mechanisms produce the same relationship between waveform shape, phase-dependent responsiveness and interregional influence remains to be tested.

The two-node model separates asymmetry in a local waveform from asymmetry in measured interactions. For identical nodes with reciprocal coupling and equal independent noise, analytical PDC is nonzero but equal in both directions, and imaginary coherency cancels even when the local waveforms are asymmetric. Unidirectional coupling breaks this exchange symmetry, producing higher PDC and nonzero imaginary coherency. Phase-locked stimulation of one node also changes the pair’s interaction measures while its structural connections remain fixed. The cancellation in the unstimulated reciprocal pair depends on identical node dynamics and inputs, and may not hold with heterogeneous nodes, unequal noise, shared input or unequal delays. In real recordings, imaginary coherency describes lagged dependence, and pairwise PDC remains susceptible to unobserved common drivers.

Temporal irreversibility captures different properties at the level of one signal and that of a network. Derivative skewness is sensitive to differences in how quickly a local waveform rises and falls. The bivariate irreversibility index instead compares transformed lagged correlations between two signals in forward and reversed time [20]. Earlier whole-brain modelling linked a related measure of non-reversibility to asymmetric effective connectivity and heterogeneous axonal delays [28]. In our simulated networks, changing synaptic rise and decay times also changed irreversibility, and this dependence persisted across the tested network sizes. The difference between directed and symmetric networks became smaller as network size increased. Network irreversibility in this model therefore reflects local kinetics as well as the pattern of connections. It is worth nothing, however, that neither derivative skewness nor the bivariate index alone measures thermodynamic entropy production or information transmission.

In the human recordings, the ascent–descent ratio was most strongly associated with local derivative skewness, whereas mean instantaneous frequency showed stronger associations with interregional measures. How much instantaneous frequency varied across the average alpha cycle, measured as the range of its phase-aligned profile divided by its mean, was also associated with interregional measures after accounting for mean frequency and the two landmark ratios. Thus, the cycle’s average rate did not fully capture its relationship with estimates of cortical interaction. These observations suggest that comparisons of cortical interactions should account for the temporal geometry of the local oscillation and also motivate testing which features best predict downstream responses. The observed associations in the present study could provide an empirical starting point for that test.

The macaque ECoG recordings showed the same negative associations between mean instantaneous frequency and the interregional measures. The human association between ascent– descent ratio and local irreversibility, however, was not statistically supported in the macaque data. Spatial summation could be a contributing factor in this difference, since it can alter waveform geometry [29], although ECoG can retain local activity [30]. The datasets also differ in clinical status, stimulation history and behavioural state, so this comparison cannot identify the cause of the discrepancy. The 31 macaque sessions came from only four animals, which limits conclusions about consistency across animals. Matched recordings across modalities and physiological conditions would help establish when the local waveform association is present.

The perturbation analysis showed a link between local kinetics and the propagation of stimulation responses. A pulse applied to the model population advances or delays its oscillation according to the phase at which it arrives, and the same local kinetics shape the waveform and the profile of this response. Coupling then determines the available routes of propagation. Thus, waveform morphology could in theory help identify temporal windows in which stimulation has a stronger downstream effect. Testing this prediction would require phase-locked stimulation across waveform states, with frequency, amplitude and stimulation intensity controlled [31]. In the human recordings, stimulation showed an effect on local waveform shape, but said changes in shape were unrelated to changes in connectivity. In addition, the stimulation was not locked to alpha phase. Cancellation of phase-specific effects is one possible explanation for this null association, but only direct testing could show the validity of this hypothesis.

The scope of the model and measurements imposes several further limits. For instance, the simulated output is a synaptic population variable. Therefore, translating it into an electrode potential requires a forward model that accounts for neuronal and recording geometry. Linear spectra and nonlinear phase responses describe different operating regimes, connected here through shared circuit parameters. Uniform time rescaling tests the role of relative time scales within that circuit, but validation across physiological rhythms will require their distinct generators to be considered. The empirical analysis focused on alpha activity, and human electrode placement reflected clinical needs in drug-resistant epilepsy.

Taken together, our findings support a physiological account of waveform morphology in the study of cortical communication. The geometry of an oscillation carries information about local temporal dynamics that complements its frequency, power and phase. Those dynamics also determine how the circuit receives and transmits input. Measuring waveform geometry alongside interregional interactions, and testing their relationship through phase-resolved perturbations, offers an experimental route from local circuit physiology to the coordination of activity across cortical networks.

## Online methods

### Modified Wilson-Cowan Model with dual-exponential synaptic kinetics

We extended the Wilson–Cowan model by passing excitatory and inhibitory population rates through separate two-stage synaptic filters before they feed back into the circuit. Independent synaptic rise and decay time constants allowed us to vary the synaptic response while keeping the fast rate time constant fixed. Recurrent nonlinearities generated oscillation harmonics, and synaptic filtering changed their relative amplitudes and phases.

### Firing rate dynamics

The classical Wilson-Cowan model is composed of one excitatory (*E*) and one inhibitory (*I*) population. Here, we first separated the instantaneous population firing rates (*R*_*E*_, *R*_*I*_) from the slow synaptic currents. In the model, the firing rates are governed by a fast membrane time constant (*τ*_*fast*_) and a sigmoidal transfer function S(*v*) that converts the net input voltage into a normalised firing rate [24]:

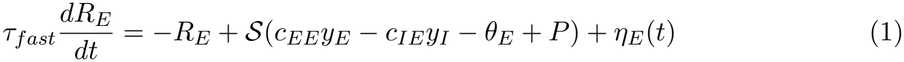

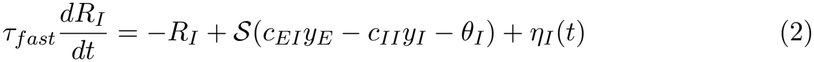

where *c*_*jk*_ is the coupling weight from population *k* to population *j, θ* is the firing threshold, *P* is the external excitatory drive, and *η*(*t*) is an additive Gaussian white noise process. The sigmoidal response function is defined as:

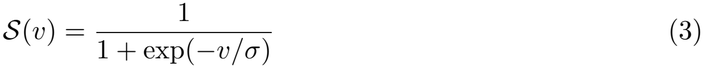

where *σ* is the steepness of the response curve.

### Dual-exponential synaptic filtering

To represent finite synaptic integration using a phenomenological kernel motivated by receptor kinetics [2], we replaced the standard single-pole low-pass filter of the Wilson-Cowan model with a double-exponential synaptic filter. This was done by cascading two first-order ordinary differential equations for each population, as follows. For the excitatory population, the firing rate *R*_*E*_ drives the synaptic rise state variable (*x*_*E*_), which in turn drives the synaptic decay state variable (*y*_*E*_):

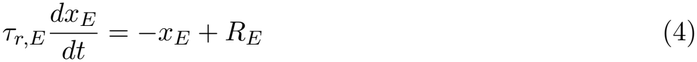

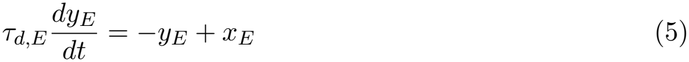

Similarly, for the inhibitory population:

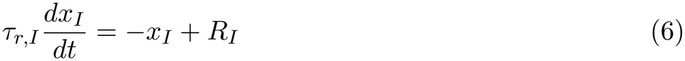

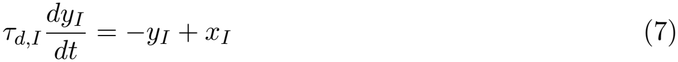

Here, *τ*_*r*_ and *τ*_*d*_ are the two kernel time scales, conventionally labelled rise and decay. The measured rise and fall durations of the emergent oscillation depend on the full circuit dynamics. The cascade is invariant to their exchange, and at equality it has the alpha-function kernel *K*(*s*) = *se*^*−s/τ*^ */τ* ^2^ for *s* ≥ 0, which remains causal and temporally asymmetric. The variable *y*_*E*_ represents the filtered population output.

By independently tuning *τ*_*r*_ and *τ*_*d*_, this model allows for controlled variation of waveform morphology, including rise–decay asymmetry and sharpness.

### Linear response and synaptic transmission

We linearised the model around its stationary operating point and tracked the eigenvalues of the local Jacobian to identify the Hopf bifurcation. The synaptic transfer function is

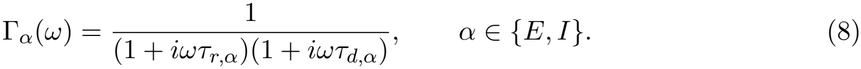

Thus each synaptic channel contributes a second-order low-pass factor. The isolated synaptic kernel is non-resonant: any resonance is generated by the closed excitatory–inhibitory feedback loop. The operating-point equations, local gains, Jacobian and reduced frequency-domain response are given in Supplementary Note S1.

The response matrix determines the power spectrum of small fluctuations around the stationary state. Writing *G*(*ω*) = *M* (*ω*)^*−*1^ for the linear propagator and *Q*(*ω*) for the spectral covariance of the inputs, the firing-rate spectral matrix is *S*_*R*_(*ω*) = *G*(*ω*)*Q*(*ω*)*G*(*ω*)^*†*^, where ^*†*^ denotes conjugate transpose. The spectra of the filtered outputs additionally include the synaptic transfer factors. We located the resonant frequency by minimising | det *M* (*ω*)|, which approximates the maximum response norm near a well-isolated resonance. The spectral derivation, explicit response matrix and resonance criterion are given in Supplementary Note S2.

### Weak coupling and phase-dependent susceptibility

We introduced weak inter-node coupling through the excitatory synaptic channel: the filtered excitatory output *y*_*E,j*_ of node *j* perturbs the excitatory rate equation of node *i* through its local gain S^*′*^(*u*_*E,i*_), with coupling strength *κ*_*ji*_. For an isolated stable limit cycle, projection onto the infinitesimal phase-response curve and averaging over one cycle yields

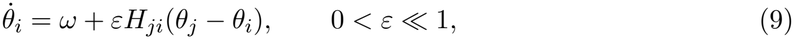

where *H*_*ji*_ is the phase interaction function. It expresses the influence of node *j* on node *i* as the overlap of postsynaptic susceptibility, local gain and the incoming waveform. Synaptic time constants shape the waveform’s temporal profile and phase lag. Together with changes in the recipient’s dynamics, these determine the phases at which input most effectively advances or delays the oscillation.

For the identical reciprocal pair, locking is controlled by the odd component of the interaction function. Its even component contributes to common frequency shifts and does not by itself establish a directional imbalance between nodes. The coupling equations, adjoint phase reduction, harmonic expansion and cross-spectral derivation are given in Supplementary Note S3.

### Directed spectral measures and symmetry

For two identical nodes with symmetric structural coupling and equal independent noise, the forward and backward contributions to the cross-spectrum cancel in its imaginary part. This cancellation relies on the exchange symmetry of the controlled pair, but heterogeneous physiology or inputs can break that symmetry. Synaptic kinetics modulate the magnitude and frequency profile of the imaginary component once exchange symmetry is broken (see Supplementary Note S3).

We used squared generalised partial directed coherence (gPDC) [27], computed from the reduced excitatory two-node response operator *A*(*ω*):

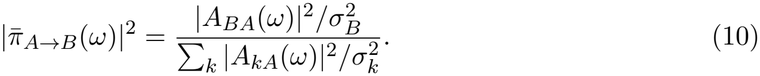

Here *A*_*ij*_ means “to *i* from *j*,” and 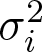 is the driving-noise variance of node *i*. Equal independent per-node noise makes this expression equal to squared classical PDC [15]. Its magnitude does not become zero in the symmetric case: the two directions are then equal, and only their difference requires directional imbalance. The full derivation and its relation to the phase interaction functions are given in Supplementary Note S3.

### Numerical simulations and validation

We validated the analytical results above by means of direct numerical simulations. Model parameters are summarised in Supplementary Table S1. Unless otherwise specified, we used a fast membrane time constant of *τ*_fast_ = 5 ms, recurrent couplings *c*_*EE*_ = 16, *c*_*IE*_ = 12, *c*_*EI*_ = 15, *c*_*II*_ = 3, a sigmoid function with steepness *σ* = 1 and thresholds of *θ*_*E*_ = 4.5 and *θ*_*I*_ = 4.0. The synaptic time constants were set to slow, theta-range values (*τ*_*r,E*_ = 3 ms, *τ*_*d,E*_ = 35 ms, *τ*_*r,I*_ = 5 ms, *τ*_*d,I*_ = 25 ms) in order for the synaptic decay time to be an appreciable fraction of the oscillation period. The stochastic differential equations were integrated with the Euler-Maruyama method with a time step of 0.1 ms. Independent additive Gaussian white noise entered the two firing-rate equations of each node.

#### Operating regimes

For each parameter set we located the Hopf bifurcation by performing a grid search over the external drive *P* and tracking the leading complex-conjugate eigenvalue pair of the Jacobian *J*. We identified the value of *P* at which its real part crossed zero, as well as its corresponding oscillation frequency. Two regimes were used: for the connectivity measures, which were derived from linear-response theory, we used a regime below the bi-furcation (small-fluctuation regime). We checked the small-fluctuation regime by comparing fluctuation amplitude with the limit-cycle amplitude. Separately, waveform morphology and both irreversibility statistics were measured above the bifurcation, in the self-oscillating limit cycle regime.

#### Waveform morphology

The waveform features were extracted from the simulated signal by means of a cycle-by-cycle approach analogous to that of Cole and Voytek [6]. Peaks and troughs were detected, and for each cycle we computed the rise time, the decay time, and the rise–decay symmetry (the rise time divided by the sum of rise and decay times).

#### Connectivity estimation

For pairs of nodes, the auto- and cross-spectra were estimated using Welch’s method. Coherence, imaginary coherency and the cross-spectral phase were obtained from these spectra. To reduce variance and bias, the spectra were averaged across an ensemble of independent realisations before obtaining the coherence. The coupling between the two nodes was introduced through the excitatory population (see Supplementary Eq. (S49)). Symmetric and directional configurations were compared at matched total coupling strength, with the directional configuration set to *κ*_*BA*_ = 0 so that only A drives B. The squared generalised partial directed coherence (gPDC^2^) was computed analytically from the off-diagonal entries of the response operator (Eq. (10)). We also fitted multivariate autoregressive models across a range of orders to the simulated two-node signals and computed gPDC from each fit.

To test whether the cancellation of imaginary coherency in the symmetric pair persists above the Hopf bifurcation, we computed band-averaged imaginary coherency for symmetric and unidirectional coupling across a range of supercritical operating points, using an excitatory synaptic decay time of 15 ms. We centred the analysis band on each operating point’s simulated peak frequency and calculated 95% confidence intervals from 2,000 bootstrap resamples of 60 realisations (Supplementary Figure S2).

#### Time-irreversibility

We quantified local temporal asymmetry using the derivative skewness [32]. Reversing the signal in time flips the sign of this measure. For a stationary reversible process with finite moments, its expected value is zero, so a reliably nonzero value provides evidence of irreversibility. The measure is sensitive to differences between rising and falling slopes. Peak and trough sharpness can differ without producing the same temporal asymmetry.

Bivariate and network time-irreversibility were quantified using the forward-versus-time-reversed shifted-correlation framework of the INSIDEOUT method [20], as also applied bivariately by G-Guzmán et al. [21]. Each signal was standardised to zero mean and unit variance. The index is the mean squared difference between forward and reversed lagged correlations after the transform 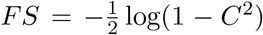, averaged over all *D*^2^ ordered channel pairs (*D* = 2 for a pair and *D* = *N* for a network). The population index disappears for a time-reversible process, so finite-sample values can be nonzero under a reversible null.

Simulation lags were defined from the isolated-node Hopf frequency *f*_H_, using *T* = round[1*/*(4*f*_H_Δ*t*_*s*_)] samples, where Δ*t*_*s*_ = 1 ms is the sampling interval after downsampling. The reference frequency was recalculated for the synaptic-kinetics sweeps and held fixed for the network-size and reciprocity sweeps. Supplementary Note S4 gives the full estimator, numerical safeguards and reference-frequency fallback rules. Supplementary Figure S3 shows the sensitivity of the measure to the lag.

The bivariate simulations were compared with independent Fourier phase-randomised surrogates, which preserve individual power spectra while randomising interchannel and within-channel harmonic phase relationships. The reference threshold was the 95th percentile of pooled individual surrogate indices (Supplementary Note S4).

Synaptic rise–decay asymmetry was varied by changing the rise–decay ratio while holding the geometric-mean synaptic time scale fixed, so that the oscillation frequency was approximately preserved.

#### Network simulations

The analysis was extended to networks of *N* nodes coupled through the excitatory channel by a weight matrix *W* . Candidate directed matrices were generated by including each possible connection independently with probability 0.5, assigning a random positive weight to each included connection and excluding self-connections. Candidates with a node receiving no directed input, or for which an exactly symmetric matrix with equal total input per node could not be constructed, were redrawn. For each accepted candidate, the symmetric condition was obtained by symmetrising and balancing the weights so that opposite-direction weights were equal and every node received the same total coupling strength. The directed condition was normalised to that same total input strength per node. Intermediate conditions interpolated between these two matrices. Network directedness is reported in Figure 3b as weighted strength reciprocity, *r* = Σ_*ij*_ min(*W*_*ij*_, *W*_*ji*_)*/* Σ_*ij*_ *W*_*ij*_ [33, 34], which equals one for an exactly symmetric matrix. Synaptic rise–decay asymmetry was varied by changing the rise– decay ratio while holding the geometric-mean synaptic time scale fixed, so that the isolated-node oscillation frequency was approximately preserved.

The external drive was set 0.4 above the estimated Hopf bifurcation of the isolated node. This estimate was recalculated for each synaptic rise–decay ratio. Each condition used 60 independent accepted network realisations. The reciprocity sweep (Figure 3b) used *N* ∈ {8, 16, 24, 32, 48}, and the synaptic-asymmetry sweep (Figure 3c) used *N* ∈ {8, 16, 32}. The network-size sweep (Figure 3d,e) varied *N* ∈ {8, 16, 24, 32, 48} with fixed synaptic parameters and the same symmetric and directed construction settings.

#### Phase-resolved perturbation

To model focal stimulation, we delivered a brief perturbation to the excitatory rate variable *R*_*E*_ of a single node and characterised its effect as a function of perturbation phase. For the single-node phase response (Figure 4b), the node was settled onto a limit cycle and, at each of 40 phases spanning one period, advanced twice from the same state: once unperturbed and once after adding *ε* = 0.05 to *R*_*E*_. The difference between the last upward mean crossings over the following three cycles was converted to a phase shift in radians and divided by *ε* to estimate the phase response *Z*(*ϕ*). Under this crossing-time convention, a positive value denotes a later crossing. This procedure was applied to fast (*τ*_*d,E*_ = 15 ms) and slow (*τ*_*d,E*_ = 40 ms) synapses, with the drive set 1.8 above each synapse’s isolated-node Hopf estimate.

For two coupled nodes (Figure 4c), paired perturbed and unperturbed trajectories were compared at each phase under unidirectional coupling and matched-total-strength reciprocal coupling. The drive was set 0.8 above the isolated-node Hopf estimate for the fast synapse. The propagated response was the root-mean-square deviation of the receiving node’s output over the cycle following the pulse. Delivering the pulse to each node yielded the two directed responses in each configuration.

For the noise-driven analysis (Figure 4d,e), the fast-synapse pair was simulated at the same drive and with the same noise intensity as the limit-cycle two-node simulations. Upward crossings of a fixed threshold in the stimulated node’s excitatory output marked successive cycles. A pulse of amplitude 0.12 was added to *R*_*E*_ once per cycle when the target fraction *ϕ* of a nominal period, calculated from the isolated-node Hopf frequency, had elapsed since the crossing. For each phase and coupling configuration, bivariate time-irreversibility was calculated at a 50-ms lag and the absolute imaginary coherency was averaged over 2–8 Hz. Each measure was averaged across 30 independent noise realisations and compared with the corresponding unperturbed and random-phase pulse baselines.

### Intracranial sEEG acquisition and preprocessing

Human stereo-EEG (sEEG) recordings were obtained from 23 patients (see Table 1 for sociode-mographic data) undergoing stereo-EEG monitoring for the presurgical evaluation of drug-resistant epilepsy at Hôpital de l’enfant Jésus - CHU de Québec - Université Laval. All patients gave written informed consent and the study was approved by the Ethics Review Board of the CHU de Québec - Université Laval (2022-5890, Québec, QC, Canada). Electrode placement was determined entirely on clinical grounds. Patients were stereotactically implanted with 10 to 18 semi-rigid multi-lead electrodes (AdTech; diameter: 0.8 mm). Brain activity was recorded from sEEG contacts, which were 2.0 mm wide and spaced 3, 4, 5, or 6 mm apart, targeting clinically defined brain structures. A 3D T1-weighted anatomical MRI (MPRAGE, 3T Siemens Trio, Siemens AG, Erlangen, Germany) was acquired pre-implantation, consisting of 160 sagittal slices with 1 mm^3^ isotropic voxels covering the entire brain. Cortical surfaces were reconstructed from the T1-weighted MRI, and the locations of the sEEG contact were identified using a post-implantation CT scan co-registered to the pre-implantation MRI (ImaGIN toolbox, https://f-tract.eu/software/imagin/).

**Table 1:** Sociodemographic characteristics of the sEEG human patients.

| Record | Biological sex | Gender | Handedness | Age | Education |
|---|---|---|---|---|---|
| S001 | Female | Female | Right | 25 | First year of master's degree |
| S002 | Female | Female | Right | 43 | Secondary 2 |
| S003 | Female | Female | Right | 18 | Secondary 4 |
| S004<sup>b</sup> | Female | Female | Right | 25 | First year of master's degree |
| S005 | Male | Male | Right | 29 | Secondary 4 |
| S006 | Male | Male | Right | 31 | DEP |
| S007 | Male | Male | Right | 26 | DEP |
| S008 | Male | Male | Right | 30 | Secondary 2 |
| S009 | Male | Male | Right | 29 | DEP |
| S010 | Male | Male | Right | 26 | CEGEP completed; Secondary 5 basic courses |
| S011 | Female | Female | Right | 34 | DEP in esthetics |
| S012 | Female | – | Right | 25 | Secondary 5 |
| S013 | Female | Male | Right | 32 | Secondary 4 |
| S014 | Male | Male | Right | 34 | Secondary 4 |
| S015<sup>a</sup> | – | Male | Right | 34 | Secondary 4 |
| S016 | Male | Male | Right | 28 | – |
| S017 | Male | Male | Right | 44 | Secondary 5 |
| S018 | Female | Female | Right | 21 | Primary school completed |
| S019 | Female | Female | Right | 36 | DEP in sales and representation |
| S020 | Female | Female | Right | 39 | Secondary school completed |
| S021 | Female | Female | Left | 49 | DEP in welding/assembly |
| S022 | Female | Female | Left | 28 | Secondary school, AS in pastry making, and DEP in cooking |
| S023 | Female | Female | Right | 25 | Secondary school completed |
| S025 | Female | Female | Left | 53 | Secondary school and DEC in administration |
| S026 | Male | Male | Right | 33 | University education; CPA in progress (part 2); bachelor's degree completed; master's degree in progress |

Age is in years. For the "Education" column, Québec qualification acronyms are retained. "–" indicates a blank source field. <sup>a</sup>S015 is the second sEEG session of S014. <sup>b</sup>S004 is the second sEEG session of S001.

The MNI coordinates of the sEEG contacts were determined using the SPM toolbox (http://www.fil.ion.ucl.ac.uk/spm/). The sEEG data were acquired with a hardware band-pass filter of 0.1–200 Hz with a sampling rate of 512 Hz (Nyquist frequency of 256 Hz) using a Natus video-sEEG monitoring system. This configuration provides a sufficient buffer to investigate the high-gamma band up to 200 Hz without significant attenuation or aliasing. Online referencing was performed using a contact located in white matter. During acquisition, signals were bandpass filtered between 0.1 and 200 Hz. sEEG contacts contaminated by pathological epileptic activity or environmental artefacts were rejected in collaboration with the clinical team. The electrical stimulation protocol follows established clinical standards and consists of bipolar stimulation delivered to SEEG contacts by the medical team during the first week of hospitalisation, using either 50-Hz trains (1–3 mA for 5 s) or 1-Hz trains (1–3 mA for 30 s). Across the cohort this yielded 5,602 stimulation runs. Two categories of recording were excluded before analysis: files acquired at 3 Hz, of which only a single example existed, and one recording whose compressed data was unreadable.

Signals were re-referenced to a bipolar montage in which each channel was formed from a pair of adjacent contacts on the same electrode shaft. Bipolar referencing was used because the difference between neighbouring contacts suppresses activity shared along the shaft, and thus limits the spatial mixing that inflates zero-lag measures of connectivity [26]. Channels were localised using each patient’s coordinate and atlas files and assigned anatomical labels according to the AAL3 atlas, then grouped into region families comprising frontal, temporal, parietal, insular, hippocampal and occipital. Amygdalar and unassigned contacts were grouped as other [35]. A channel was retained only if its own atlas label was grey matter and its two constituent contacts resolved to a single region family matching that label. Channels sharing a single contact with the stimulation pair were excluded, but the stimulation channel itself was retained, flagged and entered the models as a covariate. Recordings with fewer than five recorded channels were discarded.

For every run we extracted a pre-stimulation window spanning −30 to −2 s relative to stimulation onset, and a 28 s post-stimulation window beginning 1 s after the end of the stimulation train, which fell at 30 s for the 1 Hz protocol and at 5 s for the 50 Hz protocol. Each segment was demeaned and notch filtered at 60, 120 and 180 Hz. Slow drift was detected from the ratio of the total fitted drift to its interquartile range and removed by detrending when this ratio exceeded 0.5. Segments were rejected if they contained non-finite samples, were flat, or showed a peak-to-peak amplitude more than fifty times the interquartile range.

The replication dataset consisted of 128-channel subdural electrocorticography recorded at 1 kHz from four macaques, obtained from the Neurotycho repository [36]. Analyses were restricted to the awake, eyes-closed condition, in which the alpha rhythm was strongest, giving 31 session recordings. Each session was divided into ten non-overlapping 28 s epochs, matching the window length used for the human data.

For the macaque data, notch filters at 50, 100 and 150 Hz accounted for the 50 Hz mains supply. Bipolar channels were constructed by minimum distance matching using the published electrode coordinate maps. Consecutive channel numbering corresponded to mutual nearest neighbours for only 23% of pairs. Across sessions, this yielded 1,431 bipolar channel–session observations. Because each session contained many channel pairs, partial directed coherence was estimated on a random subset of 500 pairs per epoch, with model validity assessed over attempted pairs. Each macaque segment was linearly detrended. All later analysis steps were the same as for the human data.

### Empirical connectivity and waveform analysis of sEEG data

Testing the model in recorded data required a description of waveform shape that did not assume a sinusoidal basis. In order to study waveform morphology, conventional band-pass filtering was avoided because it can impose symmetry on the signal and separate harmonics from the fundamental that generates them [6]. We therefore isolated oscillations using masked empirical mode decomposition following Quinn and colleagues [12], which decomposes a signal into intrinsic mode functions without assuming a sinusoidal basis and preserves nonlinear waveform features.

Each segment was decomposed into six intrinsic mode functions by masked sifting, using mask frequencies of 80, 40, 20, 10, 5 and 2 Hz, a sift threshold of 10^*−*8^ and four mask phases, with monotonic piecewise cubic interpolation for the envelopes and the Rilling stopping criterion. Instantaneous phase, frequency and amplitude were obtained by the Hilbert transform with phase smoothing over three samples. All analyses used the fourth mode, which corresponds to the 10 Hz mask and captured the alpha rhythm. The median instantaneous frequency of said mode was 9.98 Hz in the human recordings and 11.5 Hz in the macaque recordings. The alpha band was chosen because it is the dominant rhythm of the cortex at rest and, like other low-frequency cortical rhythms, is known to depart from a sinusoidal waveform [5].

Individual cycles were identified from the instantaneous phase of that mode. Cycles were retained only if they were well formed, exceeded the tenth amplitude percentile of the segment, and had a maximum instantaneous frequency and an instantaneous frequency range below 20 Hz. Segments yielding fewer than five surviving cycles were excluded. For each retained cycle we computed the mean instantaneous frequency, the ascent-to-descent ratio, defined as the fraction of the cycle spent rising and equal to 0.5 for a symmetric waveform, and the peak to trough ratio, defined as the relative position of the descending zero crossing. These were averaged across cycles to give one value per channel per segment. Cycles were then projected onto a regularly sampled grid of 48 phase points to give the phase-aligned instantaneous frequency profile, which is flat for a sinusoid. From this we derived a rate-normalised descriptor of within-cycle shape, defined as the range-to-mean ratio of the phase-aligned mean instantaneous-frequency profile. This descriptor was entered into an additional model alongside mean frequency and the two cycle ratios, to test whether within-cycle shape predicts connectivity independently of mean instantaneous frequency.

***Univariate time-irreversibility*** was quantified for each channel as the skewness of the temporal derivative of the alpha mode, which changes sign under time reversal and has zero population value for a stationary reversible signal with finite moments. The signed value was retained so that the direction of the association with the signed waveform descriptors was preserved. Connectivity was estimated between every pair of bipolar channels within a recording, except pairs sharing a constituent contact (for example B1–B2 and B2–B3), whose signals are not independent. Coherence, taken as the magnitude of the coherency, and the imaginary part of the coherency were computed by the multitaper method using four-second segments. The imaginary part was summarised by its absolute value, as the sign reflects only the arbitrary ordering of the two channels. The band was 8 to 11 Hz for the human data and 8 to 13 Hz for the macaque data, the wider window being necessary because the macaque alpha mode was centred near the upper edge of the human band. Directed coupling was estimated by the squared generalised partial directed coherence [27], here obtained from a vector autoregressive model fitted to signals downsampled to 128 Hz, with model order selected up to a maximum of thirty and residual whiteness verified by the Ljung–Box test.

***Bivariate time-irreversibility*** was computed with the same forward versus time reversed shifted correlation framework used for the simulations [20], applied to the alpha mode of each pair. We used the full matrix form of the index, including the diagonal terms, so that the empirical measure is the same quantity as the one computed on the model. The time shift was set to a quarter of the dominant oscillation period, as in the simulations, but evaluated separately for each pair as *T* = round(*fs/*(4*f*)), where f is the mean instantaneous frequency of the two channels. In the simulations all nodes share a single oscillation frequency, so a quarter period is a single global shift. However, in the real data the alpha frequency varies across channels, and a fixed shift would correspond to a different fraction of the cycle at each one. Because the index is evaluated at a phase-referenced lag, holding the shift fixed would add a frequency dependence through the lag selected by the estimator.

***Pairwise connectivity*** estimates were summarised at the channel level by averaging over all pairs involving that channel. Coherence, imaginary coherency and bivariate time-irreversibility were averaged symmetrically, while partial directed coherence was summarised as the difference between a channel’s mean outgoing and mean incoming values, so that positive values indicate a net driver. For the macaque data, estimates were first averaged across the ten epochs of each session, giving one value per bipolar channel per session. For the main analysis, we included only channel observations for which we could estimate instantaneous frequency, both wave-form ratios, local time irreversibility and all four measures of interaction with other channels. This left 320,255 channel–run observations from 23 patients and 1,431 bipolar channel–session observations from 31 macaque sessions.

The primary analysis used linear mixed-effects models, one per outcome, in which the standardised outcome was regressed on the three standardised waveform features together with the available covariates, with a random intercept for the grouping variable. For the human data the covariates were brain region, stimulation protocol and stimulated-channel status, and the grouping variable was patient. For the macaque data the grouping variable was session. Models were fitted by restricted maximum likelihood using Powell’s optimiser, reverting to maximum likelihood if the random-effect variance reached its boundary. The fifteen waveform coefficients were corrected together using the Benjamini–Hochberg procedure [37]. The same three waveform predictors and five outcomes were analysed in both datasets, with the grouping and available covariates described above.

We performed three further analyses to assess robustness. To reduce dependence on channel-level pooling, the same model was fitted separately within each patient or session, and the resulting coefficients per effect were tested against zero with a one-sample Wilcoxon signed-rank test, corrected across the fifteen effects. To test the model’s predictions in the unit in which they are formulated, we also worked at the pairwise level, correlating the difference in waveform between two channels with their connectivity within each run, averaging across runs within patient, and evaluating the result against a null generated by permuting channel feature labels while holding the connectivity matrix fixed, with confidence intervals obtained by bootstrapping across patients, using 20,000 resamples. Confidence intervals on the rank correlations were likewise obtained by resampling the 23 patients with replacement, using 500 resamples, so that the interval reflects variability across patients.

Descriptive frequency associations stratified by stimulated-channel status (Supplementary Figure S8), stimulation protocol (Supplementary Figure S9) and brain region (Supplementary Figure S10) are also shown.

Changes in waveform between the pre- and post-stimulation windows were compared between stimulated and non-stimulated contacts with a two-sided Mann–Whitney *U* test.

The code used to generate all figures is available at https://github.com/pablonuneznovo/waveform_morphology/.

## Acknowledgements

This work was supported by the Canada Excellence Research Chair in Neuroplasticity. P.T. was in receipt of a NSERC Discovery Grant. This work was funded by ‘Centro de Investigación Biomédica en Red en Bioingeniería, Biomateriales y Nanomedicina (CIBER-BBN)’ through ‘Instituto de Salud Carlos III’ co-funded with ERDF funds.

