## Supplementary Information for "Waveform morphology links local circuit kinetics to cortical interactions"

#### Supplementary Notes

#### S1 Linear synaptic filtering and local Jacobian

To understand how finite synaptic kinetics shape the onset of oscillations, we first isolate the local linear dynamics around a stationary operating point. Harmonic distortion and waveform asymmetry are nonlinear effects, but they appear on top of a linear feedback structure whose poles, gains, and phase lags determine where oscillatory amplification first appears [1–4]. In this section we aim to identify the linear dynamics that will later control the weakly nonlinear regime.

The full model is defined as:

$$\tau_{\text{fast}}\dot{R}_E = -R_E + \mathcal{S}(c_{EE}y_E - c_{IE}y_I - \theta_E + P) + \eta_E(t), \tag{S1}$$

$$\tau_{\text{fast}}\dot{R}_I = -R_I + \mathcal{S}(c_{EI}y_E - c_{II}y_I - \theta_I) + \eta_I(t), \tag{S2}$$

$$\tau_{r,E}\dot{x}_E = -x_E + R_E, \tag{S3}$$

$$\tau_{d,E}\dot{y}_E = -y_E + x_E, \tag{S4}$$

$$\tau_{r,I}\dot{x}_I = -x_I + R_I, \tag{S5}$$

$$\tau_{d,I}\dot{y}_I = -y_I + x_I. \tag{S6}$$

Here $R_E$ and $R_I$ are the excitatory and inhibitory population firing rates, while $(x_\alpha, y_\alpha)$ implement a two-stage synaptic kernel with distinct rise and decay time scales. The recurrent loop is shaped by the static couplings $c_{jk}$, the slope of the gain function, and frequency-dependent synaptic filtering [1, 2, 4].

*Operating point and local gains.* Let $(R_E^*, R_I^*, x_E^*, x_I^*, y_E^*, y_I^*)$ be a stationary state. Since the synaptic subsystem is linear,

$$x_\alpha^* = R_\alpha^*, \qquad y_\alpha^* = R_\alpha^*, \tag{S7}$$

and the equilibrium is determined by

$$R_E^* = \mathcal{S}(c_{EE}R_E^* - c_{IE}R_I^* - \theta_E + P), \tag{S8}$$

$$R_I^* = \mathcal{S}(c_{EI}R_E^* - c_{II}R_I^* - \theta_I). \tag{S9}$$

Linearization about this operating point introduces the effective gains

$$g_E = \mathcal{S}'(c_{EE}R_E^* - c_{IE}R_I^* - \theta_E + P), \tag{S10}$$

$$g_I = \mathcal{S}'(c_{EI}R_E^* - c_{II}R_I^* - \theta_I), \tag{S11}$$

which renormalize the recurrent couplings. The same anatomical couplings can support different local spectra as the operating point moves along the sigmoid [2, 5].

*Jacobian.* Writing $\delta\mathbf{z} = (\delta R_E, \delta R_I, \delta x_E, \delta x_I, \delta y_E, \delta y_I)^\top$, the linearized system takes the form

$$\dot{\delta\mathbf{z}} = J\,\delta\mathbf{z} + \boldsymbol{\xi}(t), \tag{S12}$$

with

$$J = \begin{pmatrix} -\tau_{\text{fast}}^{-1} & 0 & 0 & 0 & \tau_{\text{fast}}^{-1} g_E c_{EE} & -\tau_{\text{fast}}^{-1} g_E c_{IE} \\ 0 & -\tau_{\text{fast}}^{-1} & 0 & 0 & \tau_{\text{fast}}^{-1} g_I c_{EI} & -\tau_{\text{fast}}^{-1} g_I c_{II} \\ \tau_{r,E}^{-1} & 0 & -\tau_{r,E}^{-1} & 0 & 0 & 0 \\ 0 & \tau_{r,I}^{-1} & 0 & -\tau_{r,I}^{-1} & 0 & 0 \\ 0 & 0 & \tau_{d,E}^{-1} & 0 & -\tau_{d,E}^{-1} & 0 \\ 0 & 0 & 0 & \tau_{d,I}^{-1} & 0 & -\tau_{d,I}^{-1} \end{pmatrix}.$$

The spectrum of $J$ determines the local bifurcation structure. A real eigenvalue crossing zero produces a static instability, while a complex-conjugate pair crossing the imaginary axis produces a Hopf bifurcation and hence the oscillatory branch relevant here. Hence, the Jacobian identifies the linear mode that later acquires nonlinear waveform deformation.

#### Synaptic transfer functions

We use $\Gamma_\alpha(\omega)$ for the synaptic transfer function throughout the paper, for either population $\alpha \in \{E, I\}$. The excitatory transfer function is denoted $\Gamma_E(\omega)$ (Eq. (S53)) and plotted in Figure 1e,f. The synaptic subsystem can be factored out exactly: Fourier transforming

$$\tau_{r,\alpha}\dot{x}_\alpha = -x_\alpha + R_\alpha, \tag{S13}$$

$$\tau_{d,\alpha}\dot{y}_\alpha = -y_\alpha + x_\alpha, \tag{S14}$$

gives

$$\Gamma_\alpha(\omega) = \frac{Y_\alpha(\omega)}{R_\alpha(\omega)} = \frac{1}{(1 + i\omega\tau_{r,\alpha})(1 + i\omega\tau_{d,\alpha})}, \qquad \alpha \in \{E, I\}, \tag{S15}$$

and the corresponding phase lags are

$$\varphi_\alpha(\omega) = -\arctan(\omega\tau_{r,\alpha}) - \arctan(\omega\tau_{d,\alpha}), \tag{S16}$$

Thus each synaptic channel contributes a second-order low-pass factor. In particular, the isolated synaptic kernel is non-resonant: it attenuates high frequencies and accumulates phase lag, but it does not generate a spectral peak on its own [3, 4]. Any resonance must therefore be generated by the closed excitatory-inhibitory feedback loop.

#### Reduced frequency-domain response

Eliminating the synaptic variables yields

$$(1 + i\omega\tau_{\text{fast}})R_E(\omega) = g_E\left[c_{EE}\Gamma_E(\omega)R_E(\omega) - c_{IE}\Gamma_I(\omega)R_I(\omega)\right] + \eta_E(\omega), \tag{S17}$$

$$(1 + i\omega\tau_{\text{fast}})R_I(\omega) = g_I\left[c_{EI}\Gamma_E(\omega)R_E(\omega) - c_{II}\Gamma_I(\omega)R_I(\omega)\right] + \eta_I(\omega). \tag{S18}$$

The stochastic drives $\eta_E(\omega)$ and $\eta_I(\omega)$ represent the Fourier transforms of the external fluctuations entering the excitatory and inhibitory rate equations. Equivalently,

$$M(\omega)\begin{pmatrix} R_E(\omega) \\ R_I(\omega) \end{pmatrix} = \begin{pmatrix} \eta_E(\omega) \\ \eta_I(\omega) \end{pmatrix}, \tag{S19}$$

with

$$M(\omega) = \begin{pmatrix} 1 + i\omega\tau_{\text{fast}} - g_E c_{EE}\Gamma_E(\omega) & g_E c_{IE}\Gamma_I(\omega) \\ -g_I c_{EI}\Gamma_E(\omega) & 1 + i\omega\tau_{\text{fast}} + g_I c_{II}\Gamma_I(\omega) \end{pmatrix}.$$

This reduction expresses the six-dimensional system as two populations interacting through frequency-dependent synaptic kernels. Rise and decay constants enter the feedback loop through multiplicative changes in gain and additive changes in phase delay [3, 4]. The poles of the linear response are determined by analytic continuation to the Laplace variable $s$, with $i\omega = s$:

$$\det M(-is) = 0, \qquad s \in \mathbb{C}. \tag{S20}$$

The preferred oscillatory frequency is associated with the near-singular structure of the full resolvent $M(\omega)^{-1}$. Resonant amplification occurs when recurrent E-I feedback compensates intrinsic rate relaxation and synaptic phase lags bring the loop close to constructive oscillatory closure. This linear mechanism identifies the mode from which nonlinear oscillations emerge [3, 5].

Separate rise and decay scales change the spectral gain and phase structure of the local E-I loop. These changes shift the Hopf point, move the dominant frequency and alter the linear mode from which waveform deformation develops. Retaining the synaptic cascade allows these dependencies to be examined explicitly.

#### S2 Power spectrum around the fixed point

The response matrix determines the power spectrum of small fluctuations around the stationary state. Its dependence on synaptic time constants, local gain and recurrent feedback provides access to the preferred frequency and width of the spectral peak.

*Linear response to noise.* The firing-rate fluctuations are obtained by inverting the response matrix:

$$\begin{pmatrix} R_E(\omega) \\ R_I(\omega) \end{pmatrix} = M(\omega)^{-1} \begin{pmatrix} \eta_E(\omega) \\ \eta_I(\omega) \end{pmatrix}. \tag{S21}$$

It is convenient to define the linear propagator

$$G(\omega) = M(\omega)^{-1}, \tag{S22}$$

so that

$$\mathbf{R}(\omega) = G(\omega)\,\boldsymbol{\eta}(\omega), \qquad \mathbf{R}(\omega) = \begin{pmatrix} R_E(\omega) \\ R_I(\omega) \end{pmatrix}, \quad \boldsymbol{\eta}(\omega) = \begin{pmatrix} \eta_E(\omega) \\ \eta_I(\omega) \end{pmatrix}. \tag{S23}$$

The matrix $G(\omega)$ describes how the coupled E-I populations respond to each frequency component of the noise. It combines rate relaxation, recurrent excitation and inhibition, and dual-exponential synaptic filtering.

*Noise covariance and spectral matrix.* Let the noise covariance in frequency space be

$$\langle \boldsymbol{\eta}(\omega)\boldsymbol{\eta}(\omega')^\dagger \rangle = 2\pi\,\delta(\omega - \omega')\,Q(\omega), \tag{S24}$$

where $Q(\omega)$ is the $2 \times 2$ spectral covariance matrix of the inputs, and $^\dagger$ denotes conjugate transpose. Then the spectral matrix of the firing-rate fluctuations is

$$S_R(\omega) = \langle \mathbf{R}(\omega)\mathbf{R}(\omega)^\dagger \rangle = G(\omega)\,Q(\omega)\,G(\omega)^\dagger. \tag{S25}$$

This expression holds for arbitrary stationary input covariance. For independent white-noise perturbations of equal or unequal strength,

$$Q(\omega) = \begin{pmatrix} D_E & 0 \\ 0 & D_I \end{pmatrix}, \tag{S26}$$

with constants $D_E, D_I > 0$. Then the frequency dependence of the output spectrum is entirely generated by the response matrix $G(\omega)$. The auto-spectra of the firing rates are given by the diagonal elements

$$P_{R_E}(\omega) = [S_R(\omega)]_{11}, \qquad P_{R_I}(\omega) = [S_R(\omega)]_{22}, \tag{S27}$$

while the cross-spectrum is

$$P_{R_E R_I}(\omega) = [S_R(\omega)]_{12}. \tag{S28}$$

*Spectra of synaptic variables.* To relate the rate spectra to the filtered population output, we use

$$Y_E(\omega) = \Gamma_E(\omega) R_E(\omega), \qquad Y_I(\omega) = \Gamma_I(\omega) R_I(\omega), \tag{S29}$$

the synaptic spectral matrix is obtained by an additional filtering step:

$$S_Y(\omega) = D_\Gamma(\omega)\, S_R(\omega)\, D_\Gamma(\omega)^\dagger, \tag{S30}$$

where

$$D_\Gamma(\omega) = \begin{pmatrix} \Gamma_E(\omega) & 0 \\ 0 & \Gamma_I(\omega) \end{pmatrix}. \tag{S31}$$

Hence the synaptic power spectra are

$$P_{y_E}(\omega) = |\Gamma_E(\omega)|^2 P_{R_E}(\omega), \qquad P_{y_I}(\omega) = |\Gamma_I(\omega)|^2 P_{R_I}(\omega), \tag{S32}$$

and the synaptic cross-spectrum is

$$P_{y_E y_I}(\omega) = \Gamma_E(\omega)\Gamma_I(\omega)^* P_{R_E R_I}(\omega). \tag{S33}$$

The rate spectra reflect recurrent amplification and feedback. The spectra of $y_E$ and $y_I$ additionally describe the effect of second-order synaptic filtering on those fluctuations and therefore relate the rate dynamics to the waveform of the filtered output.

#### Explicit form of the response matrix

For a $2 \times 2$ system, the inverse can be written explicitly. Define

$$a(\omega) = 1 + i\omega\tau_{\text{fast}} - g_E c_{EE} \Gamma_E(\omega), \tag{S34}$$

$$b(\omega) = g_E c_{IE} \Gamma_I(\omega), \tag{S35}$$

$$c(\omega) = -g_I c_{EI} \Gamma_E(\omega), \tag{S36}$$

$$d(\omega) = 1 + i\omega\tau_{\text{fast}} + g_I c_{II} \Gamma_I(\omega). \tag{S37}$$

Then

$$M(\omega) = \begin{pmatrix} a(\omega) & b(\omega) \\ c(\omega) & d(\omega) \end{pmatrix}, \tag{S38}$$

with determinant

$$\Delta(\omega) = \det M(\omega) = a(\omega)d(\omega) - b(\omega)c(\omega). \tag{S39}$$

Therefore,

$$G(\omega) = M(\omega)^{-1} = \frac{1}{\Delta(\omega)} \begin{pmatrix} d(\omega) & -b(\omega) \\ -c(\omega) & a(\omega) \end{pmatrix}. \tag{S40}$$

As $|\Delta(\omega)|$ approaches zero, the linear response grows and spectral power increases. This near-singularity of the feedback matrix also marks the approach to oscillatory instability in the deterministic system.

For independent white noise with covariance $Q = \mathrm{diag}(D_E, D_I)$, the firing-rate spectra become

$$P_{R_E}(\omega) = \frac{|d(\omega)|^2 D_E + |b(\omega)|^2 D_I}{|\Delta(\omega)|^2}, \tag{S41}$$

$$P_{R_I}(\omega) = \frac{|c(\omega)|^2 D_E + |a(\omega)|^2 D_I}{|\Delta(\omega)|^2}, \tag{S42}$$

$$P_{R_E R_I}(\omega) = \frac{-d(\omega)c(\omega)^* D_E - b(\omega)a(\omega)^* D_I}{|\Delta(\omega)|^2}. \tag{S43}$$

The numerators describe how the different noise sources drive each population. The common denominator describes collective amplification and, near a well-isolated resonance, largely determines the location and width of the spectral peak.

##### Peak frequency and resonance

We define the peak or resonant frequency generally as the frequency at which the response norm is maximal,

$$\omega_{\text{peak}} = \arg\max_{\omega} \|G(\omega)\|. \tag{S44}$$

Because $G(\omega) = M(\omega)^{-1} \propto 1/\Delta(\omega)$, this is closely tracked by the simpler

$$\omega_{\text{peak}} \approx \arg\min_{\omega} |\Delta(\omega)|, \tag{S45}$$

whenever the resonance is sharp enough that the $1/|\Delta(\omega)|$ singularity dominates the comparatively mild $\omega$-dependence of the numerator in Eq. (S40). The two do not need to coincide exactly away from a well-isolated resonance. We use the latter throughout, since the model operates close to the Hopf bifurcation where this condition holds.

The determinant criterion approximates the norm-based definition near a well-isolated resonance. The peak depends jointly on the fast population time scale $\tau_{\text{fast}}$, the synaptic time scales $\tau_{r,\alpha}$ and $\tau_{d,\alpha}$, the operating-point gains $g_E$ and $g_I$, and the recurrent couplings $c_{jk}$. Synaptic kinetics can therefore shift the peak through changes in both filtering and phase lag within the feedback loop.

#### S3 Weakly coupled Wilson-Cowan oscillators: synaptic kinetics, phase gating, and coupling asymmetry

We now examine two weakly coupled Wilson-Cowan oscillators to relate synaptic kinetics to phase-dependent susceptibility, cross-spectra and directed connectivity. We retain the excitatory coupling channel explicitly so that the transmitted signal can be related to the local synaptic filter.

##### Two weakly coupled Wilson-Cowan nodes

Let node $i \in \{A, B\}$ have state

$$\mathbf{x}_i = (R_{E,i}, R_{I,i}, x_{E,i}, y_{E,i}, x_{I,i}, y_{I,i})^\top,$$

with intrinsic dynamics

$$\begin{aligned}
\tau_{\text{fast}}\dot{R}_{E,i} &= -R_{E,i} + \mathcal{S}(c_{EE}y_{E,i} - c_{IE}y_{I,i} - \theta_E + P)\,, \\
\tau_{\text{fast}}\dot{R}_{I,i} &= -R_{I,i} + \mathcal{S}(c_{EI}y_{E,i} - c_{II}y_{I,i} - \theta_I)\,, \\
\tau_{r,E}\dot{x}_{E,i} &= -x_{E,i} + R_{E,i}, \qquad \tau_{d,E}\dot{y}_{E,i} = -y_{E,i} + x_{E,i}, \\
\tau_{r,I}\dot{x}_{I,i} &= -x_{I,i} + R_{I,i}, \qquad \tau_{d,I}\dot{y}_{I,i} = -y_{I,i} + x_{I,i}.
\end{aligned} \tag{S46}$$

We denote this compactly by

$$\dot{\mathbf{x}}_i = \mathbf{F}(\mathbf{x}_i). \tag{S47}$$

We then introduce weak inter-node coupling through the excitatory synaptic channel. The postsynaptic excitatory drive of node $i$ receives a small contribution from the filtered excitatory output of node $j$. The coupled system is therefore

$$\dot{\mathbf{x}}_i = \mathbf{F}(\mathbf{x}_i) + \varepsilon\,\mathbf{G}_{ji}(\mathbf{x}_j, \mathbf{x}_i), \qquad 0 < \varepsilon \ll 1, \tag{S48}$$

with

$$\mathbf{G}_{ji}(\mathbf{x}_j, \mathbf{x}_i) = \frac{\kappa_{ji}}{\tau_{\text{fast}}} \begin{pmatrix} \mathcal{S}'(u_{E,i})\,y_{E,j} \\ 0 \\ 0 \\ 0 \\ 0 \\ 0 \end{pmatrix}, \tag{S49}$$

where

$$u_{E,i} = c_{EE}y_{E,i} - c_{IE}y_{I,i} - \theta_E + P. \tag{S50}$$

The coefficient $\kappa_{ji}$ is the inter-node coupling strength. Equation (S49) makes the coupling channel explicit: the presynaptic signal is the filtered excitatory output $y_{E,j}$, and it perturbs the excitatory rate equation of the receiving node through the local gain factor $\mathcal{S}'(u_{E,i})$.

The transmitted signal $y_{E,j}$ is the output of the second-order synaptic cascade driven by the firing rate $R_{E,j}$. In the time domain,

$$y_{E,j}(t) = \int_0^\infty K_E(s)\,R_{E,j}(t-s)\,ds, \tag{S51}$$

with kernel

$$K_E(s) = \frac{e^{-s/\tau_{d,E}} - e^{-s/\tau_{r,E}}}{\tau_{d,E} - \tau_{r,E}} \qquad (\tau_{r,E} \neq \tau_{d,E}), \tag{S52}$$

and in the frequency domain,

$$\Gamma_E(\omega) = \frac{Y_E(\omega)}{R_E(\omega)} = \frac{1}{(1 + i\omega\tau_{r,E})(1 + i\omega\tau_{d,E})}. \tag{S53}$$

We use $\Gamma_\alpha(\omega)$ for synaptic transfer functions and reserve $H_{ji}(\phi)$ below for phase interaction functions. These quantities describe synaptic filtering and phase-dependent interaction, respectively.

#### Weak coupling, limit cycles, and phase reduction

Assume that for $\varepsilon = 0$ each isolated node admits a stable limit cycle

$$\mathbf{x}_i(t) = \mathbf{X}(\theta_i), \qquad \dot{\theta}_i = \omega, \tag{S54}$$

with period $T = 2\pi/\omega$. The oscillatory regime considered here is precisely the one generated by the Hopf bifurcation discussed above. Weak coupling then perturbs the motion only slightly away from the cycle. Transverse amplitude perturbations relax on the intrinsic timescale of the local oscillator, whereas phase perturbations accumulate on the slower coupling timescale $O(\varepsilon^{-1})$. This separation justifies a phase description.

Let $\mathbf{Z}(\theta)$ be the infinitesimal phase-response curve, i.e. the periodic adjoint satisfying

$$\frac{d\mathbf{Z}}{d\theta} = -\frac{1}{\omega}\left[D\mathbf{F}(\mathbf{X}(\theta))\right]^{\top}\mathbf{Z}(\theta), \qquad \mathbf{Z}(\theta)\cdot\mathbf{F}(\mathbf{X}(\theta)) = \omega. \tag{S55}$$

Projecting the perturbation onto the tangent phase direction gives, to first order in $\varepsilon$,

$$\dot{\theta}_i = \omega + \varepsilon\,\mathbf{Z}(\theta_i)\cdot\mathbf{G}_{ji}\big(\mathbf{X}(\theta_j),\mathbf{X}(\theta_i)\big). \tag{S56}$$

Averaging over one cycle yields the reduced phase equation

$$\dot{\theta}_i = \omega + \varepsilon\,H_{ji}(\theta_j - \theta_i), \tag{S57}$$

where the phase interaction function is

$$H_{ji}(\phi) = \frac{1}{2\pi}\int_0^{2\pi}\mathbf{Z}(\vartheta)\cdot\mathbf{G}_{ji}\big(\mathbf{X}(\vartheta+\phi),\mathbf{X}(\vartheta)\big)\,d\vartheta. \tag{S58}$$

For the Wilson-Cowan coupling in Eq. (S49), this becomes

$$H_{ji}(\phi) = \frac{\kappa_{ji}}{2\pi\,\tau_{\text{fast}}}\int_0^{2\pi} Z_E(\vartheta)\,\mathcal{S}'\big(u_E(\vartheta)\big)\,y_E(\vartheta+\phi)\,d\vartheta, \tag{S59}$$

where $Z_E$ is the PRC component associated with the excitatory rate equation and $u_E(\vartheta) = c_{EE}y_E(\vartheta) - c_{IE}y_I(\vartheta) - \theta_E + P$ is the excitatory input along the uncoupled cycle.

Equation (S59) expresses the influence of node $j$ on node $i$ as the overlap of three phase-dependent quantities: postsynaptic susceptibility $Z_E(\vartheta)$, local gain $\mathcal{S}'(u_E(\vartheta))$, and the incoming waveform $y_E(\vartheta+\phi)$. Synaptic time constants shape the waveform's temporal profile and phase lag. Together with changes in the recipient's dynamics, these determine the phases at which input most effectively advances or delays the oscillation.

The phase difference $\psi = \theta_B - \theta_A$ then evolves according to

$$\dot{\psi} = \Delta\omega + \varepsilon\big[H_{AB}(-\psi) - H_{BA}(\psi)\big], \tag{S60}$$

with $\Delta\omega = \omega_B - \omega_A$. In the symmetric case $H_{AB} = H_{BA} = H$,

$$\dot{\psi} = \Delta\omega - 2\varepsilon H_{\text{odd}}(\psi), \qquad H_{\text{odd}}(\psi) = \frac{H(\psi) - H(-\psi)}{2}. \tag{S61}$$

Thus, for the identical reciprocal pair, locking is controlled by the odd component of the interaction function. Its even component contributes to common frequency shifts and does not by itself establish a directional imbalance between nodes.

#### From synaptic filtering to phase interactions and cross-spectra

The phase-interaction kernel and the frequency-domain response both depend on the synaptic transfer function evaluated at the harmonics of the oscillation.

As shown above, the synaptic output of the sending node admits a harmonic expansion

$$y_E(t) = \sum_{n\geq 1} A_n^{(y)} \cos(n\omega t + \varphi_n), \tag{S62}$$

with coefficients determined by the synaptic filter,

$$A_n^{(y)} e^{i\varphi_n} = \Gamma_E(n\omega)\, A_n^{(R)} e^{i\chi_n}. \tag{S63}$$

Substituting this expansion into the interaction formula in Eq. (S59) yields

$$H_{ji}(\phi) = \sum_{n\geq 1} \left[a_n^{(ji)} \sin(n\phi) + b_n^{(ji)} \cos(n\phi)\right], \tag{S64}$$

with coefficients that scale as

$$a_n^{(ji)},\, b_n^{(ji)} \propto \kappa_{ji}\, \Gamma_E(n\omega)\, A_n^{(R)}\, \mathcal{Z}_n. \tag{S65}$$

This expression makes the qualitative dependence explicit, written schematically as:

$$\Gamma_E(n\omega) \implies \text{harmonic content of } H_{ji}(\phi). \tag{S66}$$

The interaction kernel projects the synaptically filtered coupling onto the oscillatory harmonics of the limit cycle. The same transfer function appears in the linear response of the coupled system. In the frequency domain, inter-node coupling enters through the off-diagonal operator

$$\mathcal{K}_{BA}(\omega) \propto \kappa_{AB}\, \Gamma_E(\omega), \tag{S67}$$

so that the cross-spectrum satisfies

$$S_{AB}(\omega) = \left[\mathcal{R}(\omega)\, \Sigma_\xi(\omega)\, \mathcal{R}^\dagger(\omega)\right]_{AB}. \tag{S68}$$

Near the oscillatory regime, where power is concentrated around harmonics of $\omega$, one can isolate the dominant contribution and write schematically

$$S_{AB}(n\omega) \propto Q_{AB}(n\omega)\, \Gamma_E(n\omega), \tag{S69}$$

with $Q_{AB}$ collecting local gains and noise structure. This gives the second, likewise schematic, link in the mapping:

$$\Gamma_E(n\omega) \implies S_{AB}(n\omega). \tag{S70}$$

Equations (S66)-(S70) summarize the shared dependencies of the phase and spectral descriptions in the following schematic:

$$\Gamma_E(n\omega) \implies H_{ji}(\phi) \implies S_{AB}(\omega). \tag{S71}$$

The phase interaction function and cross-spectrum describe complementary consequences of synaptic filtering. The phase-interaction function describes susceptibility along the limit cycle, while the cross-spectrum describes the frequency-resolved relationship between the populations. Both are governed by the same harmonic weighting imposed by $\Gamma_E(\omega)$. Their shared dependence on synaptic kinetics connects changes in phase locking, coherence and spectral structure. Adjusting $\tau_r$ and $\tau_d$ reshapes $\Gamma_E(\omega)$, which in turn reweights harmonic contributions to $H(\phi)$ and redistributes spectral power and phase relations in $S_{AB}(\omega)$.

#### Synaptic rise-decay asymmetry, phase bias, and imaginary coherence

We now turn to the role of synaptic rise-decay asymmetry. For finite synaptic time constants, the transfer function is complex-valued and introduces a frequency-dependent phase lag,

$$\Gamma_E(\omega) = |\Gamma_E(\omega)|\, e^{i\varphi_E(\omega)}. \tag{S72}$$

Finite synaptic time constants produce a phase lag for both equal and unequal rise and decay scales. Varying their ratio changes the frequency profile of the lag and the weighting of harmonics. These changes affect the phase-interaction kernel and, when exchange symmetry is broken, the imaginary component of the cross-spectrum.

At the level of the interaction function, the complex phase of $\Gamma_E(n\omega)$ shifts each harmonic contribution,

$$H_{ji}(\phi) \sim \sum_{n\geq 1} |\Gamma_E(n\omega)| A_n \sin\big(n\phi + \varphi_E(n\omega) + \psi_n\big), \tag{S73}$$

where $\psi_n$ collects contributions from the PRC and local gain. Because $\varphi_E(n\omega)$ depends on $n$, the resulting kernel is generally not antisymmetric. This produces a nonzero even component,

$$H_{\text{even}}(\phi) = \frac{H(\phi) + H(-\phi)}{2}, \tag{S74}$$

and hence a phase-dependent bias in how perturbations are transmitted across the cycle. Synaptic kinetics therefore reshape phase-dependent transmission. This does not by itself generate a directional imbalance between identical reciprocally coupled nodes.

The full cross-spectrum combines the forward and backward pathways. Under the assumptions of the identical-node pair, the two contributions represented schematically in Eq. (S69) enter as a complex-conjugate pair. Writing $h(\omega)$ for the local closed-loop excitatory response of an isolated node, and retaining terms to first order in the coupling, the cross-spectrum of a pair driven by independent noise of equal intensity $D$ is

$$S_{AB}(\omega) = D\, |h(\omega)|^2 \left[\kappa_{BA}\, h(\omega)\, \Gamma_E(\omega) + \kappa_{AB}\, h^*(\omega)\, \Gamma_E^*(\omega)\right], \tag{S75}$$

in which the channel from $A$ to $B$ and the channel from $B$ to $A$ are complex conjugates of one another. Two consequences follow. For a structurally symmetric pair, $\kappa_{AB} = \kappa_{BA} = \kappa$, the bracket reduces to $2\kappa\, \Re[h(\omega)\Gamma_E(\omega)]$, which is real, so that

$$\Im\, S_{AB}(\omega) = 0 \qquad (\kappa_{AB} = \kappa_{BA}), \tag{S76}$$

irrespective of the synaptic phase $\varphi_E(\omega)$: the synaptic lag cannot, on its own, produce an imaginary coherence. In general,

$$\Im\, S_{AB}(\omega) \propto (\kappa_{BA} - \kappa_{AB})\, \Im[h(\omega)\, \Gamma_E(\omega)], \tag{S77}$$

so that the imaginary part is gated by the coupling asymmetry $\kappa_{BA} - \kappa_{AB}$ and only then shaped by the synaptic phase, which enters through $\Im[h\Gamma_E]$. For this controlled pair, imaginary coherence requires directional coupling, and synaptic rise-decay asymmetry modulates its magnitude and frequency profile (Figure 2). This cancellation is exact under the idealised assumptions adopted for the derivation: two identical nodes, symmetric structural coupling, equal and independent per-node noise, no common input, and no heterogeneity between them. Any departure from these, unequal noise, common input, heterogeneous nodes, delays, or referencing effects, can reintroduce a nonzero imaginary component even without directional coupling. The network case in the main Methods is one such departure, since heterogeneous neighbourhoods break the pairwise exchange symmetry the two-node result relies on.

The relationship to directed connectivity depends on the symmetry of the full system. Measures such as PDC, imaginary coherence and the bivariate arrow of time all depend on asymmetries in the coupling operator itself, for example $\kappa_{AB} \neq \kappa_{BA}$. Imaginary coherence and the bivariate arrow of time vanish for a structurally symmetric, identically driven pair, and PDC loses its directional asymmetry under the same condition. Synaptic rise-decay asymmetry modulates the magnitude and frequency profile of these measures through the frequency-dependent phase of $\Gamma_E(\omega)$. Synaptic rise-decay asymmetry therefore reshapes, in a frequency-dependent manner, the directionality that is carried by the coupling, and this is reflected in both phase-domain and frequency-domain observables. We make this distinction precise numerically in the Results, where perturbation responses also demonstrate phase-selective influence.

Hence, synaptic rise-decay asymmetry acts as a phase-selective filter. Through its complex transfer function, it shapes the harmonic structure of the interaction kernel, induces phase bias in oscillator coupling, and modulates the imaginary component available in the cross-spectrum once exchange symmetry is broken by coupling asymmetry. These effects are different manifestations of a single mechanism: the frequency-dependent phase lag introduced by the synaptic dynamics.

#### Directed interactions and partial directed coherence

For the analytical PDC calculation, we use a reduced excitatory two-node response. Let $\mathbf{X}(\omega) = (R_{E,A}(\omega), R_{E,B}(\omega))^\top$ and let $\boldsymbol{\xi}(\omega)$ denote the corresponding independent driving fluctuations. We write

$$\mathbf{X}(\omega) = A(\omega)^{-1}\boldsymbol{\xi}(\omega), \tag{S78}$$

where the operator used for the analytical calculation is

$$A(\omega) = \begin{pmatrix} a_0(\omega) & -g_E\kappa_{BA}\Gamma_E(\omega) \\ -g_E\kappa_{AB}\Gamma_E(\omega) & a_0(\omega) \end{pmatrix}, \qquad a_0(\omega) = 1 + i\omega\tau_{\text{fast}} - g_E c_{EE}\Gamma_E(\omega). \tag{S79}$$

Here $\kappa_{AB}$ denotes coupling from node $A$ to node $B$, with the weak-coupling scale absorbed into $\kappa_{AB}$ and $\kappa_{BA}$. Thus $A_{BA}(\omega) = -g_E\kappa_{AB}\Gamma_E(\omega)$ is the entry for $A \to B$, with the reverse direction given analogously. The diagonal term retains excitatory rate relaxation and excitatory self-feedback. This reduced operator was used for analytical PDC. The full local excitatory–inhibitory response is described by $M(\omega)$ in Supplementary Note S1.

A standard directed measure is partial directed coherence. We use the generalised definition of Baccalá, Sameshima & Takahashi [6], which normalises by each node's own driving-noise variance $\sigma_i^2$ so that the measure remains scale-invariant across channels with different intrinsic variability:

$$|\bar{\pi}_{A\to B}(\omega)|^2 = \frac{|A_{BA}(\omega)|^2/\sigma_B^2}{\sum_k |A_{kA}(\omega)|^2/\sigma_k^2}, \tag{S80}$$

with analogous expressions for the reverse direction. In the present model every node receives independent noise of the same fixed intensity $D$ regardless of coupling configuration, so $\sigma_A^2 = \sigma_B^2$ identically. The variance-normalisation therefore cancels exactly, and Eq. (S80) reduces to the square of the classical (unweighted) partial directed coherence, $\pi_{A\to B}(\omega) = |A_{BA}(\omega)|/\sqrt{\sum_k |A_{kA}(\omega)|^2}$, of Baccalá & Sameshima [7]. This equivalence requires the equal, independent per-node noise injection used in the model. Equation (S79) shows that synaptic kinetics affect PDC directly through the same transfer factor that shaped the interaction kernel and the cross-spectrum. The effect of synaptic filtering can be examined separately from the imbalance between the two coupling directions. Both a nonzero imaginary coherence and a directional imbalance in PDC require asymmetry in the effective coupling operator, for example $\kappa_{AB} \neq \kappa_{BA}$ (Eq. (S77)). What is specific to PDC is that its *magnitude* need not vanish in the symmetric case: the two directions are then equal, $\bar{\pi}_{A\to B} = \bar{\pi}_{B\to A}$, and it is only their *difference* that requires directed coupling. Synaptic kinetics enter all of these measures through the same transfer factor $\Gamma_E(\omega)$, and modulate their magnitude and frequency profile under the stated symmetry conditions.

At the phase-reduced level, an analogous distinction appears in the interaction kernels. If

$$H_{AB}(\phi) \neq H_{BA}(-\phi), \tag{S81}$$

the phase drift acquires a directional bias. Even components in $H(\phi)$ reflect asymmetry of phase-dependent transmission and susceptibility and do not by themselves imply a causal asymmetry in the multivariate linear sense captured by PDC. Both descriptions depend on the same synaptically filtered coupling channel. Phase reduction emphasizes how that channel gates perturbations across the oscillatory cycle, while spectral measures emphasize how the same channel shapes cross-spectra and directed transfer.

#### S4 Bivariate and network irreversibility estimator

Bivariate and network time-irreversibility were quantified using the forward-versus-time-reversed shifted-correlation framework of the INSIDEOUT method [8], as also applied bivariately by G-Guzmán et al. [9]. For a set of $D$ signals $x_i(t)$ ($i = 1, \dots, D$. $D = 2$ for the bivariate case, $D = N$ for the network case), each signal was standardized to zero mean and unit variance over the full analysed record. Let $x_i^{(r)}(t)$ denote the corresponding time-reversed signal. The forward and reversed time-shifted correlation matrices at lag $T$ are

$$C^{\mathrm{f}}_{ij}(T) = \langle x_i(t)\, x_j(t+T)\rangle_t\,, \tag{S82}$$

$$C^{\mathrm{r}}_{ij}(T) = \left\langle x_i^{(r)}(t)\, x_j^{(r)}(t+T)\right\rangle_t\,, \tag{S83}$$

where the averages are calculated over the overlapping samples at each lag. Before applying the logarithmic transform, correlation estimates were clipped to $[-0.999999, 0.999999]$ to avoid singularities. Each correlation was then converted to a Gaussian mutual-information-like quantity,

$$FS^{\mathrm{f}}_{ij}(T) = -\tfrac{1}{2}\log\!\big(1 - [C^{\mathrm{f}}_{ij}(T)]^2\big), \qquad FS^{\mathrm{r}}_{ij}(T) = -\tfrac{1}{2}\log\!\big(1 - [C^{\mathrm{r}}_{ij}(T)]^2\big). \tag{S84}$$

The irreversibility index is the mean squared difference between the forward and reversed transformed correlations,

$$I(T) = \frac{1}{D^2}\sum_{i=1}^{D}\sum_{j=1}^{D}\big(FS^{\mathrm{f}}_{ij}(T) - FS^{\mathrm{r}}_{ij}(T)\big)^2. \tag{S85}$$

The population index vanishes for a time-reversible process. Positive values indicate asymmetry in the transformed lagged correlations, although finite-sample estimates can be nonzero under a reversible null. The average includes all $D^2$ ordered channel pairs. For this estimator, the reversed correlation matrix is the transpose of the forward matrix, so the diagonal contributions vanish up to numerical precision. For $D = 2$, the full-matrix index therefore equals one half of the off-diagonal mean. We applied Eq. (S85) uniformly to the bivariate and network cases. For networks, we additionally restricted the average to channel pairs with a direct structural connection.

Simulation lags were defined from the isolated-node Hopf frequency $f_{\mathrm{H}}$, using $T = \mathrm{round}[1/(4 f_{\mathrm{H}} \Delta t_s)]$ samples, where $\Delta t_s = 1$ ms is the sampling interval after downsampling. The reference frequency was recalculated for the synaptic-kinetics sweeps and held fixed for the network-size and reciprocity sweeps. The implementation used fallback reference frequencies when the bifurcation search failed its acceptance criteria and imposed a lower frequency bound of 1 Hz when calculating the lag. This lag defines the time scale at which the statistic was evaluated.

The bivariate simulations were compared with surrogates obtained by independent Fourier phase randomization of each channel (Figure 2f). This preserves each channel's power spectrum while randomizing relative Fourier phases across channels and the phase relationships between harmonics within each channel. The surrogate comparison therefore tests against a restricted null for the joint dynamics. Observed indices were averaged across runs, whereas the reference threshold was the 95th percentile of pooled individual surrogate indices. This threshold characterizes the pooled surrogate distribution and does not constitute a calibrated significance test of the ensemble mean.

Sensitivity to the selected lag is shown in Supplementary Figure S3.

#### Supplementary Tables

Table S1: **Model parameters used throughout, unless stated otherwise.** The fixed point is independent of the timescales. Uniform rescaling of all timescales leaves the dimensionless dynamics invariant and simply rescales the oscillation frequency. Changes in the timescales' relative values can shift the Hopf condition itself.

| Symbol | Meaning | Value |
|---|---|---|
| $\tau_{\text{fast}}$ | fast population timescale | 5 ms |
| $\tau_{r,E},\ \tau_{d,E}$ | excitatory synaptic rise, decay | 3 ms, 35 ms |
| $\tau_{r,I},\ \tau_{d,I}$ | inhibitory synaptic rise, decay | 5 ms, 25 ms |
| $c_{EE},\ c_{IE},\ c_{EI},\ c_{II}$ | recurrent coupling weights | 16, 12, 15, 3 |
| $\sigma$ | sigmoid steepness | 1.0 |
| $\theta_E,\ \theta_I$ | firing thresholds | 4.5, 4.0 |
| $P$ | external drive (Hopf at) | $\approx 1.76$ |
| $f_{\text{Hopf}}$ | emergent frequency at Hopf | $\approx 4.3$ Hz |
| $dt$ | integration step (Euler–Maruyama) | $10^{-4}$ s |
| $\kappa$ | two-node coupling (directed) | 0.5 |
| density, $\kappa_{\text{tot}}$ | network density, per-node incoming coupling | 0.5, 0.5 |

### Supplementary Figures

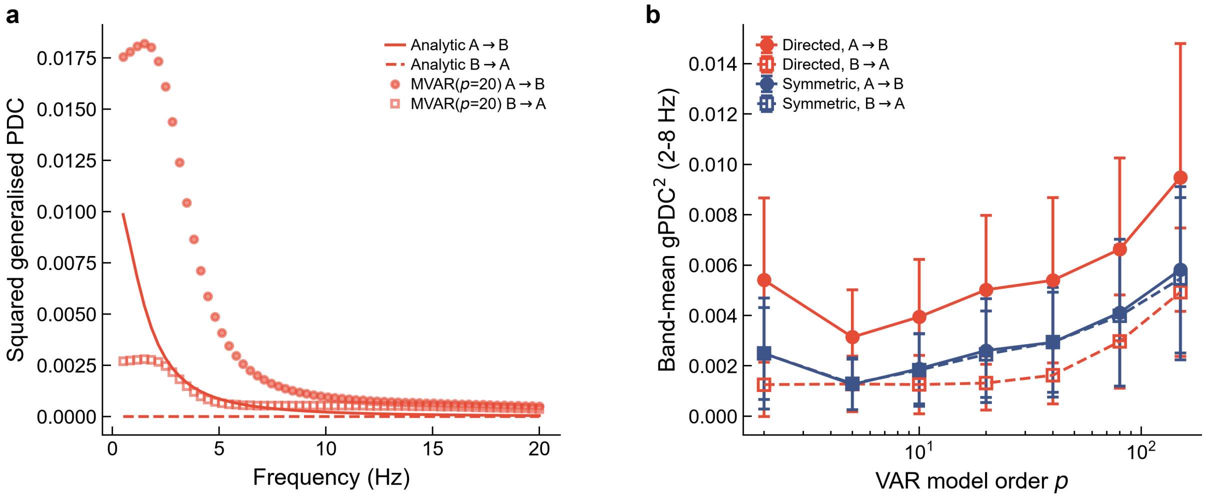


Figure S1: **Estimated squared generalised partial directed coherence depends on autoregressive model order.** gPDC$^2$ was estimated by fitting multivariate autoregressive (MVAR) models to the two-node signals from Figure 2, for directed ($A \to B$) and symmetric coupling. (a) At order $p = 20$, the estimated $A \to B$ gPDC$^2$ (points) follows the analytic prediction (solid line) in shape but not magnitude. The analytically zero $B \to A$ estimate (open squares) is instead nonzero and converges toward $A \to B$ at higher frequencies. (b) Band-averaged (2–8 Hz) gPDC$^2$ versus model order (mean ± s.d., 30 realisations).

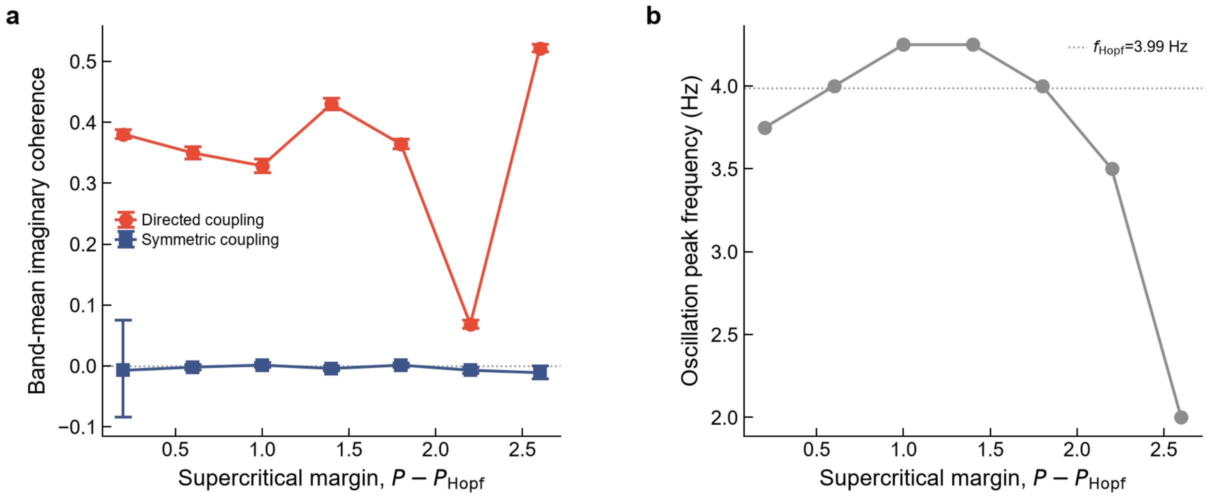


Figure S2: **Imaginary coherence remains small under symmetric coupling in the nonlinear regime.** (a) Band-mean imaginary coherence for symmetric ($\kappa_{AB} = \kappa_{BA}$) and directed ($\kappa_{BA} = 0$) coupling across supercritical margins $P - P_{\text{Hopf}}$ (fast/skewed synapse, $\tau_d = 15$ ms, as in Figure 4), with the analysis band re-centred on each margin's peak frequency (panel b). Points are bootstrap means with 95% confidence intervals (2000 resamples, 60 realisations). (b) Oscillation peak frequency at each margin.

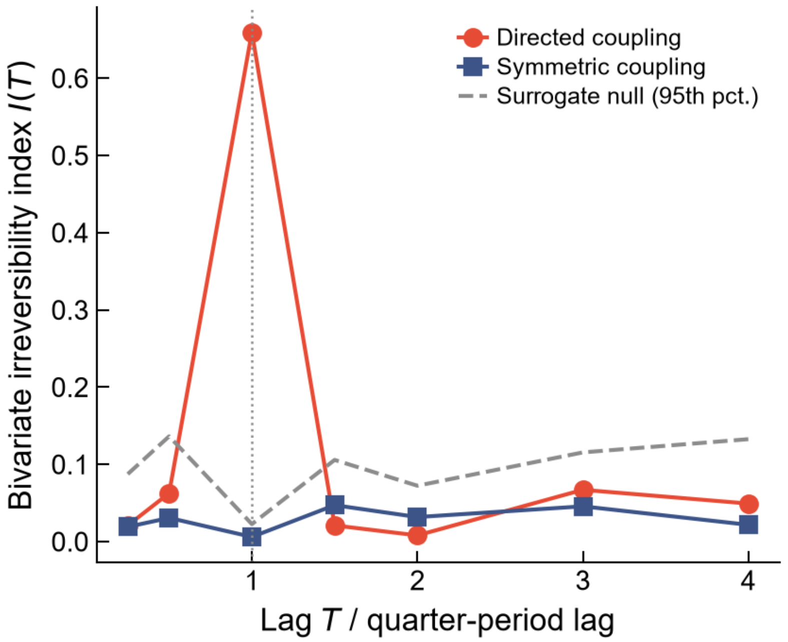


Figure S3: **Bivariate time-irreversibility depends on the selected lag.** Bivariate irreversibility $I(T)$ for directed (red circles) and symmetric (blue squares) coupling, with the 95th percentile of the pooled phase-randomised surrogate indices (grey dashed line), as a function of lag $T$. The lag is expressed relative to the quarter-period reference used for the simulations. Simulations used $\tau_d = 30$ ms and a supercritical margin $P - P_{\text{Hopf}} = 0.8$.

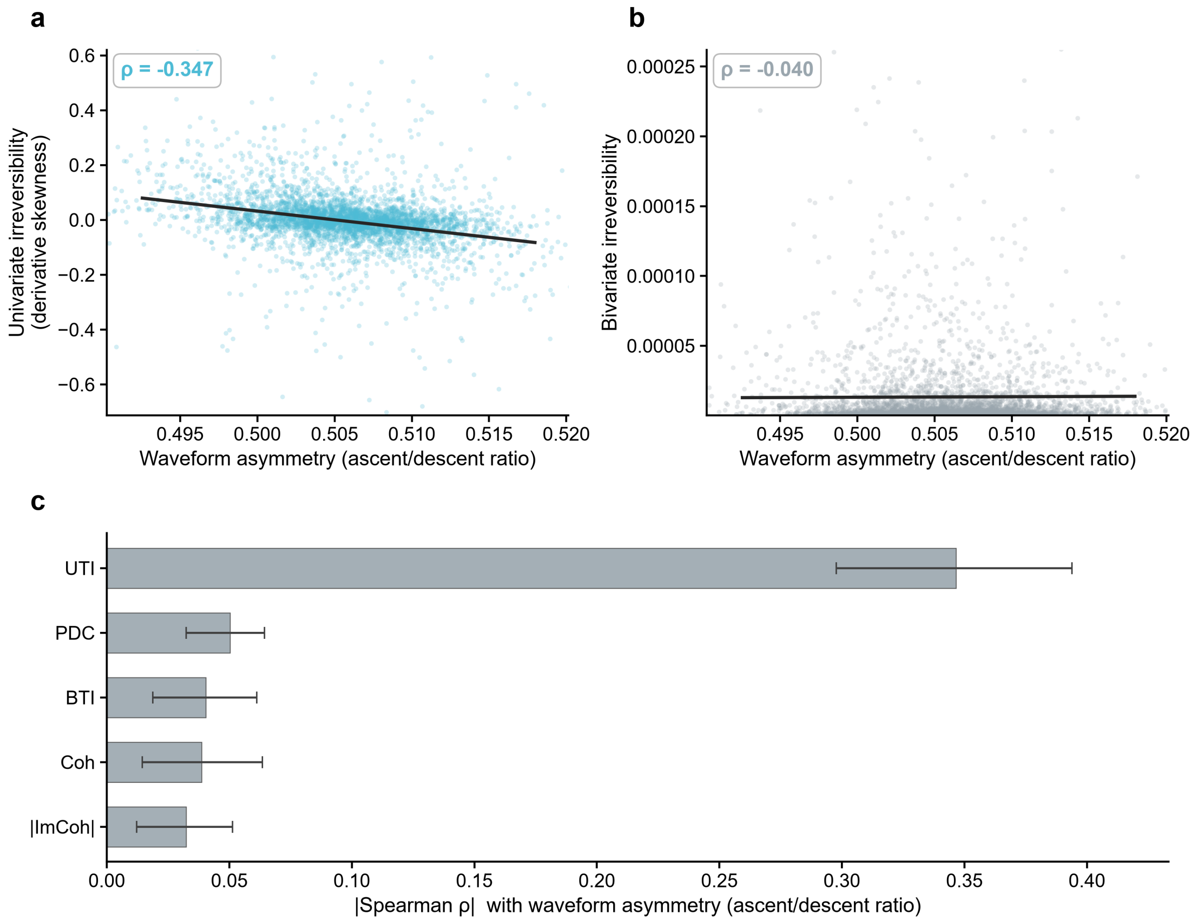


Figure S4: **Waveform asymmetry is more strongly associated with local than inter-regional temporal asymmetry.** (a) Univariate time-irreversibility against waveform asymmetry across electrodes. (b) Bivariate time-irreversibility against the same variable. Points are a random sample of 4,000 channel-level observations for legibility. The fitted line and Spearman $\rho$ inset use all 320,255 observations. (c) Absolute Spearman correlation with waveform asymmetry for all five measures, ordered by effect size. Bars show the pooled across-electrode statistic, and whiskers show the 95% confidence interval obtained by resampling the 23 patients with replacement using 500 resamples. Coh, coherence. |ImCoh|, magnitude of imaginary coherency. PDC, partial directed coherence asymmetry. BTI, bivariate time-irreversibility. UTI, univariate time-irreversibility.

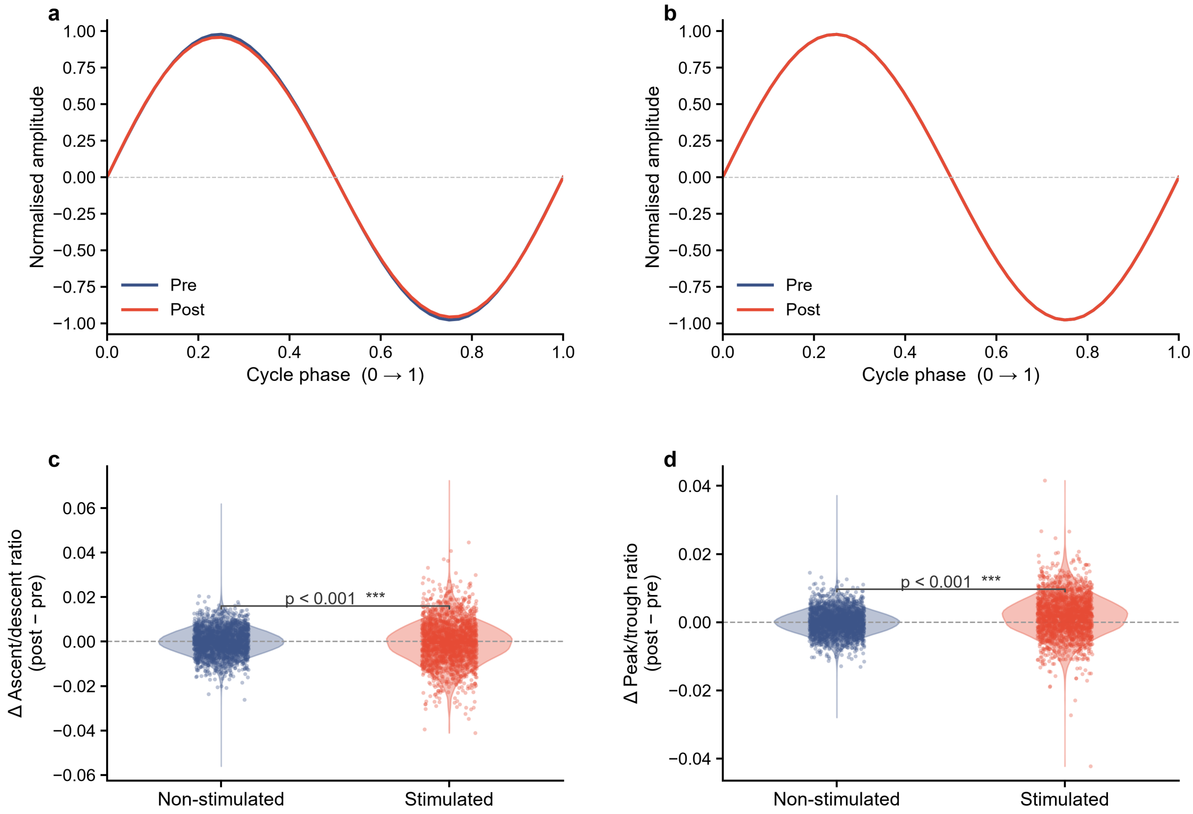


Figure S5: **Waveform changes after stimulation differ between stimulated and non-stimulated contacts.** (a) Grand-mean normalised cycle before and after stimulation at stimulated contacts, shaded ± SEM (standard error of the mean). (b) The same for non-stimulated contacts, where the average cycle is unchanged. (c,d) Changes in the ascent-to-descent and peak-to-trough ratios from the pre-stimulation to the post-stimulation window for non-stimulated and stimulated contacts. Violin plots show the distributions, and points represent individual electrodes. The reported $p$-values are from two-sided Mann–Whitney $U$ tests comparing the two groups.

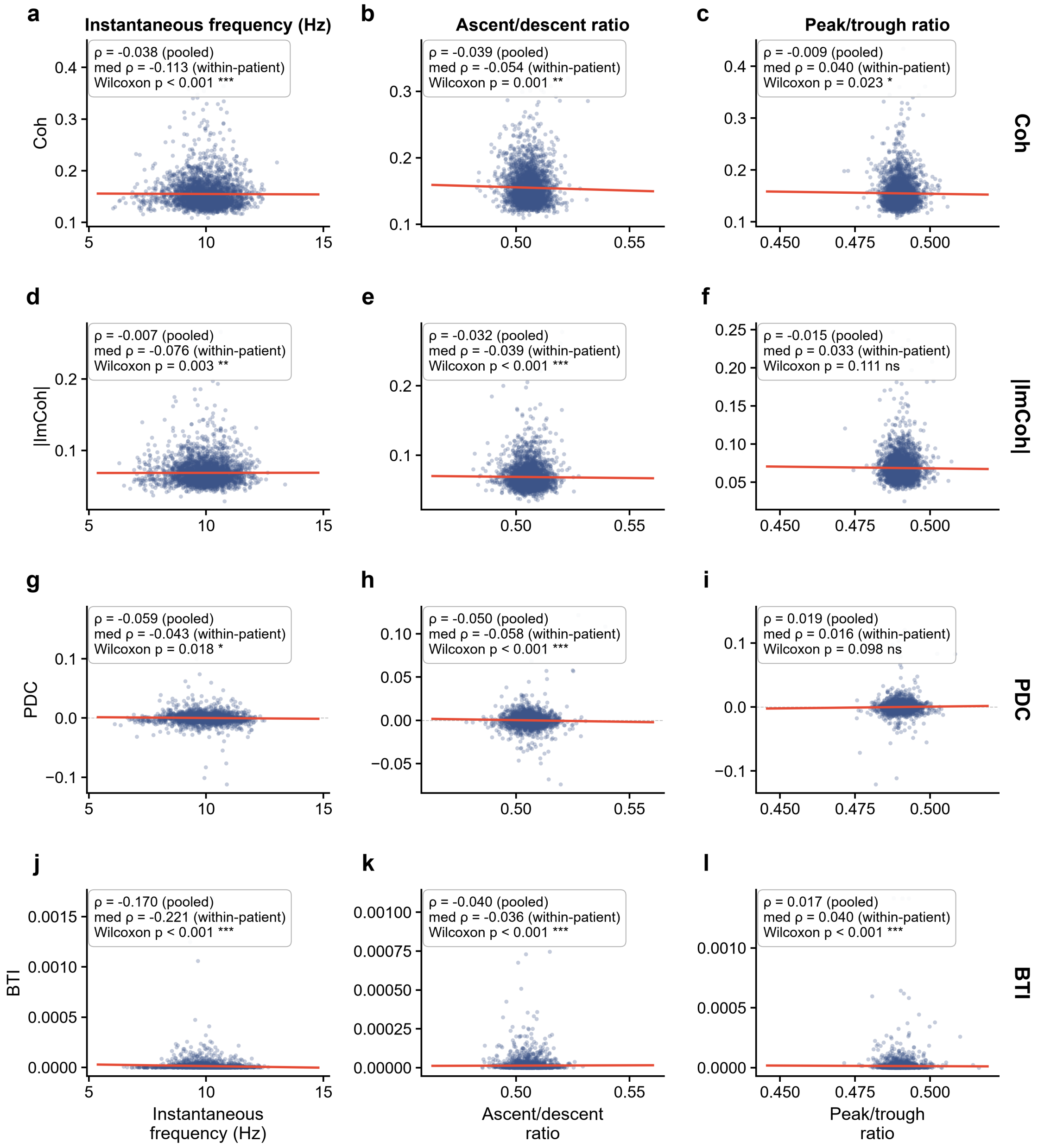


Figure S6: **Unadjusted associations between waveform features and connectivity.** (a–l) Each of the four between-channel measures (rows) is plotted against each of the three waveform features (columns) across channel-level observations. Points show a random subsample for legibility, but the fitted lines use all electrodes. Insets report the pooled Spearman $\rho$ across all electrodes, the median within-patient $\rho$, and a one-sample Wilcoxon signed-rank test of the 23 within-patient correlations. Comparison with the primary analysis shows that several associations, particularly the ordering among the between-channel measures, appeared only after brain region, stimulation protocol and patient were accounted for. Coh, coherence. |ImCoh|, magnitude of imaginary coherency. PDC, partial directed coherence asymmetry. BTI, bivariate time-irreversibility.

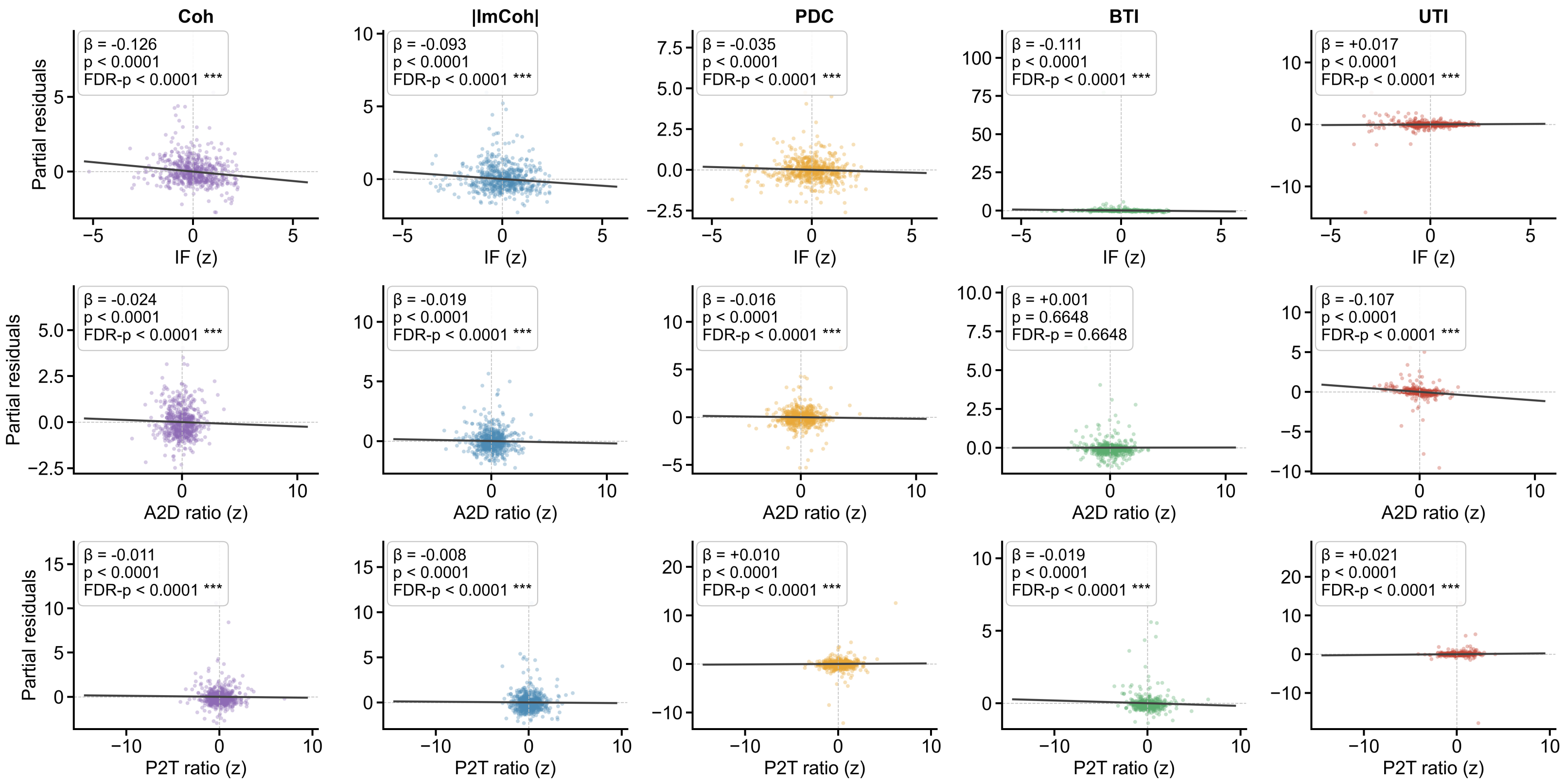


Figure S7: **Partial regressions for the primary mixed-effects models.** Each waveform feature (rows) is plotted against each outcome (columns), with the contributions of the other predictors and covariates removed. The horizontal axis shows the standardised waveform feature, and the vertical axis shows the model residuals plus the contribution of that predictor. The fitted-line slope is the standardised coefficient reported in the primary analysis. Points show a random subsample of electrodes. Stars indicate FDR-corrected significance: *** $p < 0.001$, ** $p < 0.01$, * $p < 0.05$; ns, not significant. Coh, coherence. |ImCoh|, magnitude of imaginary coherency. PDC, partial directed coherence asymmetry. BTI, bivariate time-irreversibility. UTI, univariate time-irreversibility.

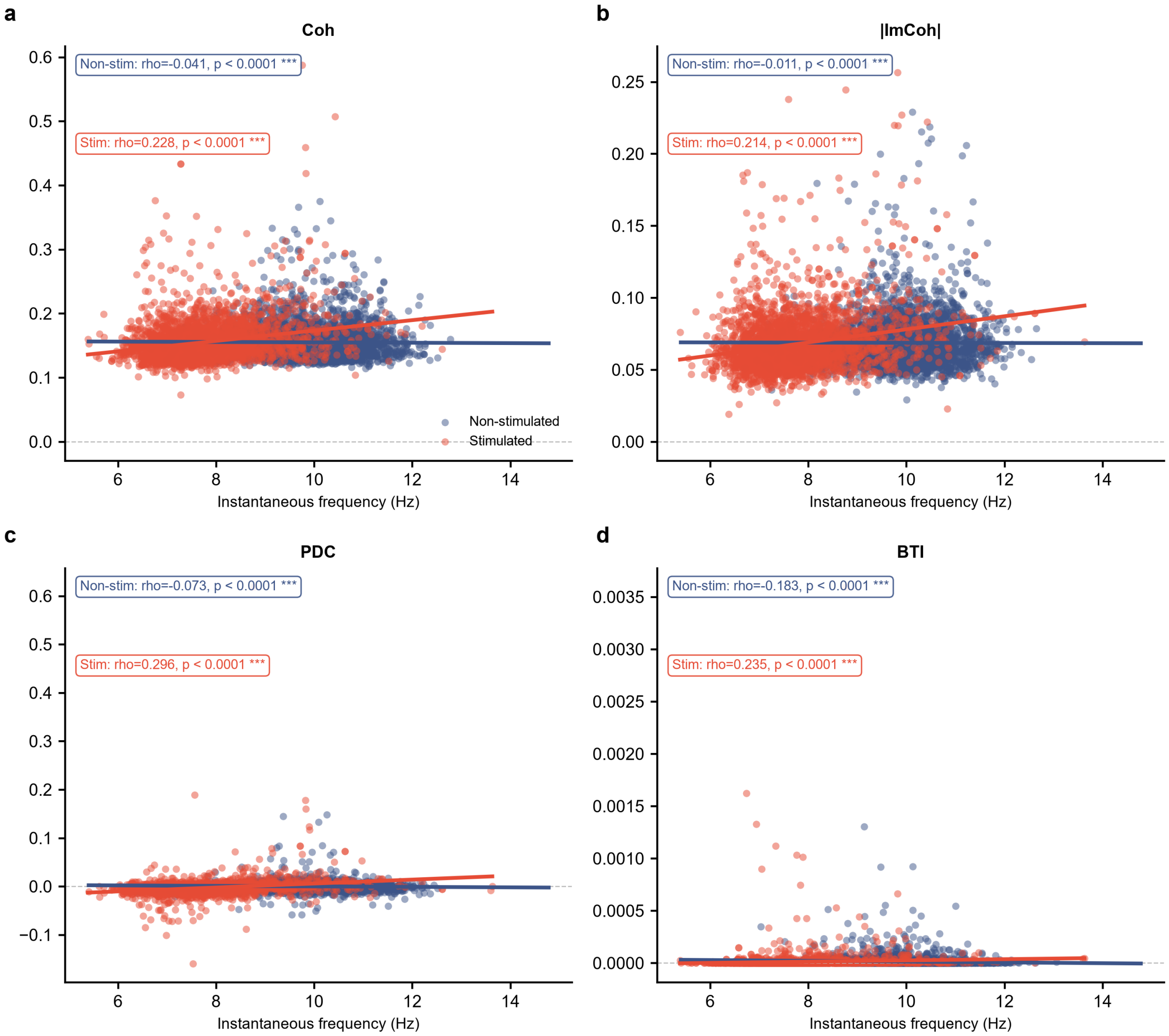


Figure S8: **Frequency associations differ between stimulated and non-stimulated contacts.** (a–d) Coherence, imaginary coherence, PDC asymmetry and bivariate time-irreversibility are plotted against instantaneous frequency, separately for stimulated (red) and non-stimulated (blue) contacts. Lines show fitted associations and insets report Spearman $\rho$ and the corresponding $p$-value for each group. The relationship is negative in non-stimulated contacts and positive in stimulated contacts for every measure. Coh, coherence. |ImCoh|, magnitude of imaginary coherency. PDC, partial directed coherence asymmetry. BTI, bivariate time-irreversibility.

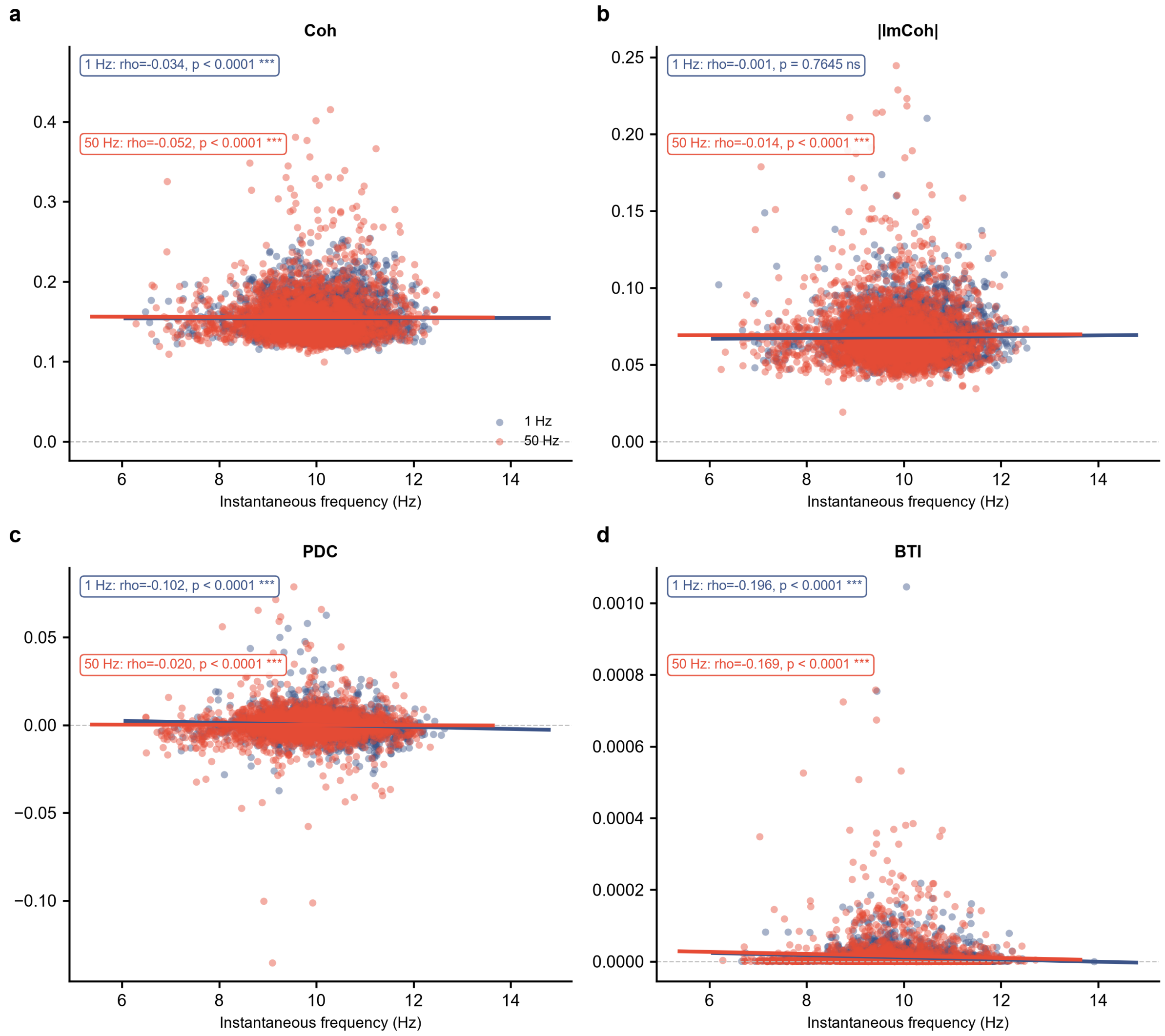


Figure S9: **Frequency associations under the two stimulation protocols.** (a–d) Associations between instantaneous frequency and the four between-channel measures are shown separately for the 1-Hz (blue) and 50-Hz (red) stimulation protocols, following the format of Supplementary Figure S8.

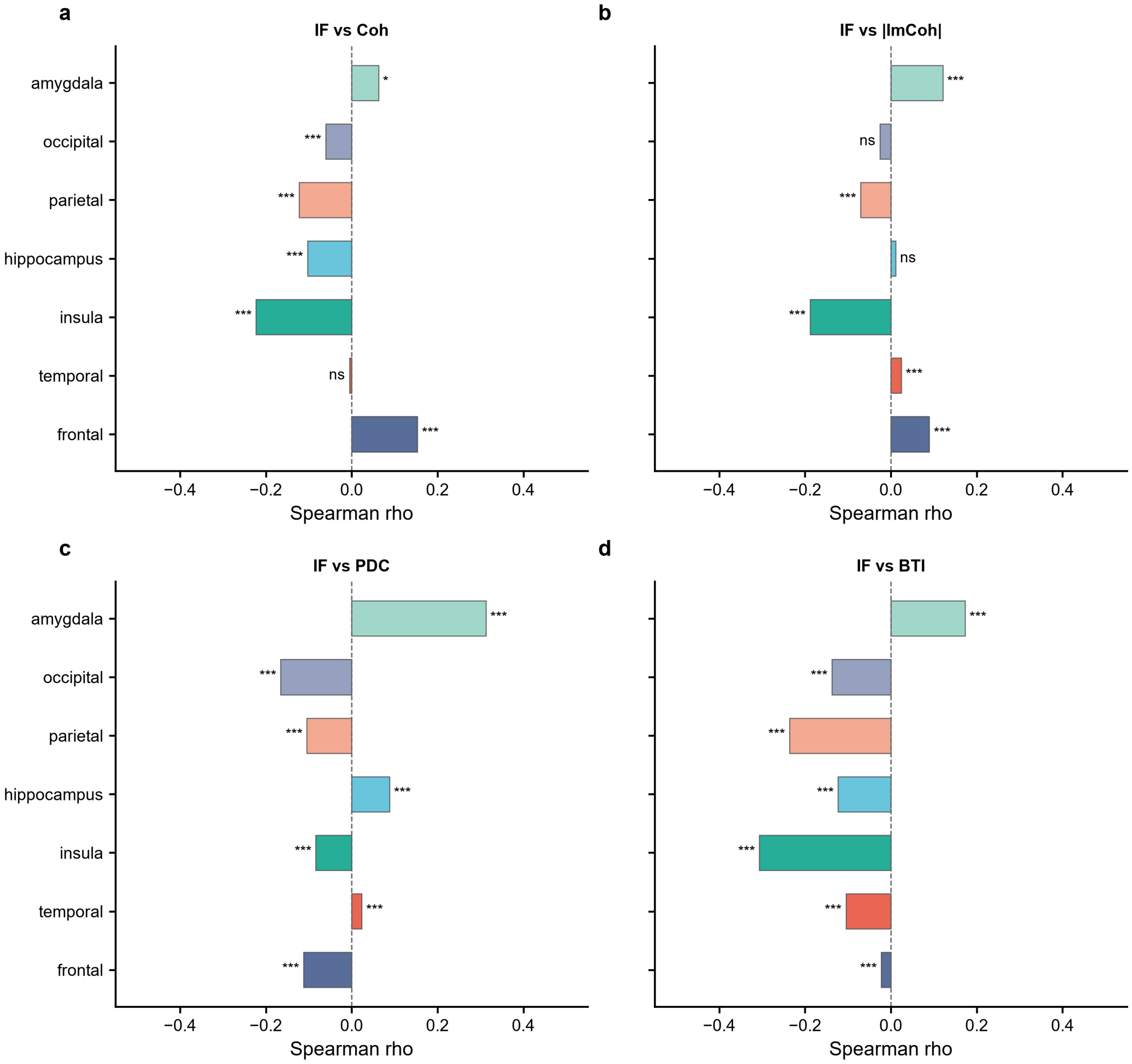


Figure S10: **Frequency associations vary across brain regions.** (a–d) Spearman correlations between instantaneous frequency and coherence, imaginary coherence, PDC asymmetry and bivariate time-irreversibility, calculated separately within each anatomical region family. Stars indicate FDR-corrected significance: *** $p < 0.001$, ** $p < 0.01$, * $p < 0.05$; ns, not significant. Coh, coherence. |ImCoh|, magnitude of imaginary coherency. PDC, partial directed coherence asymmetry. BTI, bivariate time-irreversibility.
